# Transient alterations in nucleosome distribution and sensitivity to nuclease define the THP-1 monocyte to macrophage transition

**DOI:** 10.1101/2025.02.11.637703

**Authors:** Jane M Benoit, Brandon D Buck, Mahdi Khadem, Hank W Bass, Jonathan H Dennis

## Abstract

The monocyte to macrophage transition is marked by alterations to both the structure and function of the genome, including changes in histone post-translational modifications, DNA methylation, 3D nuclear architecture, and expression of lineage specific genes. The nucleosome is the fundamental organizational unit of the eukaryotic genome and underpins both genome structure and function. However, nucleosome dynamics at promoters, which are essential for transcriptional regulation, are understudied in cellular differentiation. We conducted high-resolution chromatin structure profiling at promoters in the THP-1 cell line at eight different time points spanning PMA-induced monocyte to macrophage differentiation. We found that fewer than 10% of nucleosomes within promoters were redistributed during differentiation and only a subset of these were associated with immediate transcriptional alterations. Nucleosomes within the promoters of PMA-responsive genes were strongly positioned prior to differentiation and experienced minimal alterations during differentiation thus implying the existence of a pre-differentiation primed chromatin state. Additionally, we observed pronounced alterations in nucleosome sensitivity to MNase digestion within one hour of PMA-induced differentiation and the emergence of a highly resistant phenotype in fully differentiated cells. We found that resistance is correlated with active chromatin marks, transcription factor binding, gene expression, and higher order chromatin structure demonstrating that it is a useful measure of both genome structure and function. Together this suggests that, unlike more stable nucleosome distribution, transient sensitivity alterations may underpin new genomic functions in differentiating cells. Our results offer a framework for understanding how chromatin structural alterations potentiate cellular differentiation in a monocyte model and use methodology that is widely applicable to other systems.

**Summary sentence:** Nucleosome distribution is largely static during PMA induced monocyte differentiation while nucleosome sensitivity is highly dynamic and is associated with gene expression, active chromatin marks, transcription factor binding, and higher order chromatin structure.

## Introduction

The remarkable plasticity of myeloid progenitor cells enables multicellular organisms to mount appropriate immune responses against a diverse array of pathogens. Circulating monocytes can differentiate into effector macrophages when stimulated by macrophage colony-stimulating factor ^1,2^. Additional signals from pathogens and cytokines, such as LPS, γ-IFN, or TNFα, further refine this differentiation, leading to the generation of polarized macrophages with distinct pro-inflammatory or anti-inflammatory roles ^3,4^. Monocyte differentiation entails profound phenotypic changes as these cells extravasate into tissues, become highly phagocytic, produce reactive oxygen and nitrogen species, upregulate cytokine production, and transition from having a lobulated to a rounded nucleus ^2^. Despite the critical importance of macrophages in both innate and adaptive immune responses, the precise regulatory programs governing macrophage differentiation and activation remain incompletely understood.

Notably, monocytes and macrophages, despite sharing the same genomic DNA, exhibit substantial phenotypic and functional differences which are regulated by epigenomic modifications. These modifications play a pivotal role in the monocyte-to-macrophage transition, manifesting in widespread changes in both global chromatin interactions and local chromatin structure during differentiation. Unlike other types of cells undergoing differentiation, the terminal monocyte to macrophage transition does not involve progression through mitosis, necessitating epigenomic modifications to rapidly reprogram vast numbers of genes in a highly specific and temporarily controlled manner. Rapid DNA demethylation, mediated by the 5mC hydroxylase TET2, occurs in the first 18 hours of differentiation ^5^ and is localized in promoters, exons, and transcription termination sites ^6^. Macrophages exhibit a distinctive histone post-translational modification repertoire, characterized by increased H3K27ac and H3K4me1 marks and decreased H3K27me3 at CpG demethylated enhancer like regions ^5^ but these modifications were not always associated with immediate alterations in transcription ^7^. Chromatin remodeling during monocyte differentiation makes key regulatory regions accessible for specific transcription factor binding which allows macrophages to respond rapidly to pathogens ^8,9^. Binding of macrophage specific regulatory factors such as AP-1, EGR2, IRF4, PU.1, and STAT family members is increased at newly demethylated DNA regions during differentiation ^6^ and a subset of regulatory factors, such as MAFb, are necessary for differentiation ^10^. While the alterations in higher order chromatin structure in differentiated macrophages may be cell model specific ^11^, neither CTCF nor cohesin are required for macrophage differentiation ^12,13^. Additionally, dynamic chromatin looping facilitates the expression of key macrophage genes by connecting loci with distal AP-1 binding sites during the differentiation process ^14^. These, and other, epigenomic modifications poise promoters and enhancers in the macrophage genome to respond to pathogens and drive a macrophage specific gene regulatory program.

Eukaryotic genome regulation is primarily established and maintained through *cis* and *trans* interactions between gene promoters, regulatory elements, and regulatory factors, which interact to tune gene expression. These interactions occur in the context of highly compacted chromatin composed of individual nucleosome subunits made of ∼147bp of DNA wrapped around a histone octamer and linked together by short stretches of linker DNA ^15^. The location and density of nucleosomes can modulate regulation by their effects on underlying promoter DNA sequence elements, making them more or less accessible to their cognate regulatory factors ^16–18^. Genomes are organized in a non-random manner and specific promoters are enriched for cell-type specific regulatory factors. Our knowledge of chromatin structure and epigenomic mark dynamics comes from a number of techniques developed to map epigenetic marks, regulatory factor binding, and chromatin accessibility including ChIP, CUT&RUN, CUT&TAG, MNase-seq, ATAC-seq, FAIRE-seq, DNAse-seq, while related methods such as 3C, HiC, and derivatives have measured long-range chromatin interactions ^19,20–22,23^. However, each of these methods alone offers an incomplete understanding of genome regulation as they provide information about a single feature of the chromatin landscape and an integrated approach is needed to understand the relationship between chromatin architecture and gene regulation. Micrococcal nuclease (MNase) based assays allow for combined analyses of nucleosome positioning, nucleosome sensitivity, and cistrome occupancy which contribute to understanding the chromatin landscape on a nucleosomal level and its relation to gene regulation. ^24–30,31,3232–31^ ^30,33,34^. These assays provide a broader survey of chromatin features than is captured by a single pulldown or 3C based technique. MNase sensitivity in particular provides insight into both local and higher order chromatin structure and is correlated with gene expression and epigenomic marks (reviewed in ^35^).

Epigenetic modifications are known to be essential for the monocyte to macrophage transition but the precise chronology of these alterations during differentiation is not clear. Most epigenetic studies compare cells before and after the transition, but typically lack analysis of well-defined timepoints during the differentiation process. Temporal resolution is needed to determine the sequential order of chromatin modifications. In order to create a time-resolved epigenomic picture of the monocyte to macrophage transition, we utilized the model THP-1 monocytic cell line, which can be differentiated into macrophages by treatment with phorbol 12-myristate 13-acetate (PMA). THP-1 is a well-established immortalized model cell line to investigate cellular differentiation and innate immune responses ^36^ ^37^. THP-1-derived macrophages express near normal levels of Toll and NOD-like receptors, and stimulation with lipopolysaccharide results in strong cytokine production and phagocytic activity ^38^. These cells have a stable genetic background and can be readily and reproducibly expanded ^36,39^ which makes them an invaluable tool for studying myeloid biology. However, these cells do differ from primary human monocytes in terms of gene expression, polarization ability, and chromatin structure ^11,40,41^ which necessitate consideration when using these cells as a model for monocyte differentiation.

To clarify our understanding of promoter architecture during the monocyte to macrophage transition, we used our previously described mTSS-seq method ^42,43^ to map MNase-protected nucleosomes at high resolution across the promoters of 21,857 pol II human genes during the monocyte to macrophage transition in THP-1 cells. THP-1 monocytes were treated with PMA and samples were collected at eight time points spanning the monocyte to macrophage transition. We found that nucleosome distribution within promoters is largely stable during differentiation and that the promoter architecture of PMA-responsive genes is pre-established in undifferentiated cells. We also observed widespread alterations in sensitivity to nuclease digestion with a more resistant phenotype emerging in fully differentiated cells. Our results characterize a functional framework using nucleosome sensitivity assays to gain deeper insight into nucleosome dynamics and how they impact chromatin states during monocyte differentiation.

## Methods

### Tissue culture and nuclei isolation

THP-1 monocytes were obtained from the ATCC (TIB-202) and were cultured in RPMI-1640 media (Gibco) supplemented with 10% fetal bovine serum and 1% penicillin-streptomycin. Cells were maintained at 37°C with 5% CO_2_. Cells were exposed to 25nM phorbol 12-myristate 13-acetate (Sigma-Aldrich) for 48 hours and recovered in complete medium for 24 hours following differentiation^37^. Both adherent and suspension cells were harvested at the 1-hour, 4-hour, 8-hour, 12-hour, and 24-hour timepoints. Only adherent cells were harvested at the 48-hour and 72-hour time points. At each time point, cells were crosslinked in 1% formaldehyde (Sigma-Aldrich) and incubated for 10 minutes at room temperature while rocking. Excess formaldehyde was quenched with 125mM glycine and cells were washed twice in PBS. Cells were resuspended in nuclei isolation buffer (10 mM HEPES at pH 7.8, 2 mM MgOAc_2_, 0.3 M sucrose, 1 mM CaCl_2_ and 1% Triton-X) and pelleted by centrifugation at 1000 × *g* for 10 min at 4 °C. Nuclei were washed twice in nuclei isolation buffer and nuclei purity was confirmed by DAPI staining.

### MNase cleavage and preparation of DNA libraries

All nuclei samples were processed in replicate. 2.5×10^6^ nuclei were resuspended in 500μl nuclei isolation buffer (10 mM HEPES at pH 7.8, 2 mM MgOAc_2_, 0.3 M sucrose, 1 mM CaCl_2_ and 1% Triton X) and digested with Micrococcal nuclease (MNase) (Worthington) under light (5 U) or or heavy (200 U) conditions for 10 minutes at 37 °C. MNase reactions were halted with 50 mM EDTA. Protein-DNA crosslinks were reversed by treating the MNase-digested nuclei with 0.2 mg/mL proteinase K and 1% sodium dodecyl sulfate, and incubating overnight at 65 °C. DNA was isolated using guanidinium-isopropanol and washed with ethanol on a silica column. Mononucleosomal and sub-mononucleosomal DNA was isolated from a 2% agarose gel slice and purified by glycogen precipitation.

MNase sequencing libraries were prepared for each replicate using the Accel-NGS®2S Plus DNA Library Kit (Swift) for Illumina® using 10ng of input DNA from each digestion. Libraries were indexed with the Accel-NGS 2S Unique Dual Indexing Kit and normalized with Normalase. Library size and quality was verified with a D1000 Tapestation. Molar concentration of each indexed library was determined by KAPA quantitative PCR and size corrected using sizing information from the Tapestation.

### Solution-based sequence capture and Illumina sequencing

We used a previously validated custom designed Roche Nimblegen SeqCap EZ Library SR to capture 2000 bp regions flanking the Transcription Start Site (TSS) for every pol II gene in the human genome ^42,43^ based on the Reference Sequence (RefSeq) gene set. This approach increases sequencing depth and resolution at gene promoters by reducing the sequenced genome from ∼3.4 GB to ∼40 MB.The TSS sequences were repeat masked using human Cot-1 DNA and library adapters were masked with universal blocking oligos. Biotinylated sequence capture oligos were hybridized to input libraries for 48 hours and purified with streptavidin beads. Enriched sequences were amplified in a 15 cycle PCR amplification using TruSeq primers. Final library size and quality was validated with the Tapestation and KAPA quantitative PCR was used to determine final molar concentrations. Paired end 50bp sequences were generated on the FSU College of Medicine’s NovaSeq 6000 Illumina sequencer.

### Data processing

Raw sequencing reads were demultiplexed using Casava and adapter sequences and low quality reads were removed with Trimmomatic (version 0.39)^44^. Bowtie2 was used to align the sequences to the T2T reference human genome with the --end-to-end --no-mixed --no-discordant flags ^45^. Mitochondrial reads were filtered with samtools (version 1.10) ^46^ and duplicate reads were removed with picardtools (version 2.27.3) (Picard Toolkit. 2019. Broad Institute, GitHub Repository. https://broadinstitute.github.io/picard/; Broad Institute). Nucleosome footprints for light and heavy digests were calculated and quantile normalized with DANPOS (version 3.1.1) ^47^. The log_2_ ratio of light to heavy digests was calculated for each time point with deeptools (version 3.5.1)^48^ to determine nucleosome sensitivity to MNase digestion. Figures showing individual loci were generated with the PlotGardener package (version 1.4.1)^49^ in R version (version 4.3.2). Gene ontology was generated using the Reactome 2024 database ^50,51^ through the Enrichr webtool ^52–54^. Nucleosome occupancy and sensitivity files were uploaded to the UCSC genome browser and NCBI GEO database under accession number GSE254274.

Data for additional cell lines was processed in parallel. The HEK293 and A549 cells are from unpublished Dennis lab datasets. The GM12878 and MCF10A nucleosome data were previously published as GSE139224 and GSE134297, respectively.

External data from GEO dataset GSE96800 ^14^ were obtained and concurrently processed for comparative analysis as follows. RNA-seq data was trimmed with Trimmomatic, aligned to the T2T reference genome with HiSat2, and further processed with Stringtie and Ballgown to call differentially expressed genes ^55–57^. ChIPseq samples from GSE96800 ^14^ and GSE143981 ^58^ and CUT&RUN samples from GSE172296 ^59^ were trimmed with trimmomatic and aligned to the T2T reference genome with Bowtie2. Mitochondrial and duplicate reads were filtered with samtools (version 1.10) ^46^ and picardtools (version 2.27.3) (Picard Toolkit. 2019. Broad Institute, GitHub Repository. https://broadinstitute.github.io/picard/; Broad Institute) and peaks were called with MACS3 ^60^. HiC contact maps were viewed with Juicebox ^61^ and HPTM tracks were added from the ENCODE portal ^62,63^ (https://www.encodeproject.org/) with the following identifiers: ENCFF526GZE and ENCFF727JUN.

## Results

### Nucleosome profiles are largely stable across promoters during PMA induced monocyte differentiation

We used MNase based chromatin profiling to determine how PMA induced differentiation impacts promoter nucleosome distribution over time as described in Figure 1. We treated THP-1 monocytes with phorbol 12-myristate 13-acetate (PMA) to induced differentiation and collected samples at eight time points throughout the monocyte differentiation process at 0hr, 1hr, 4hr, 8hr, 12hr, 24hr, 48hr, and 72hr. These time points were selected to span the complete differentiation spectrum and a 24hr rest period to permit cellular recovery (as recommended by ^37^). We digested crosslinked chromatin from each time point with contrasting nuclease levels to produce light and heavy digests (Fig. 2A) which were used to generate maps of nucleosome distribution and sensitivity to nuclease (Fig. 1) (described in ^28,30^).

**Figure 1.**
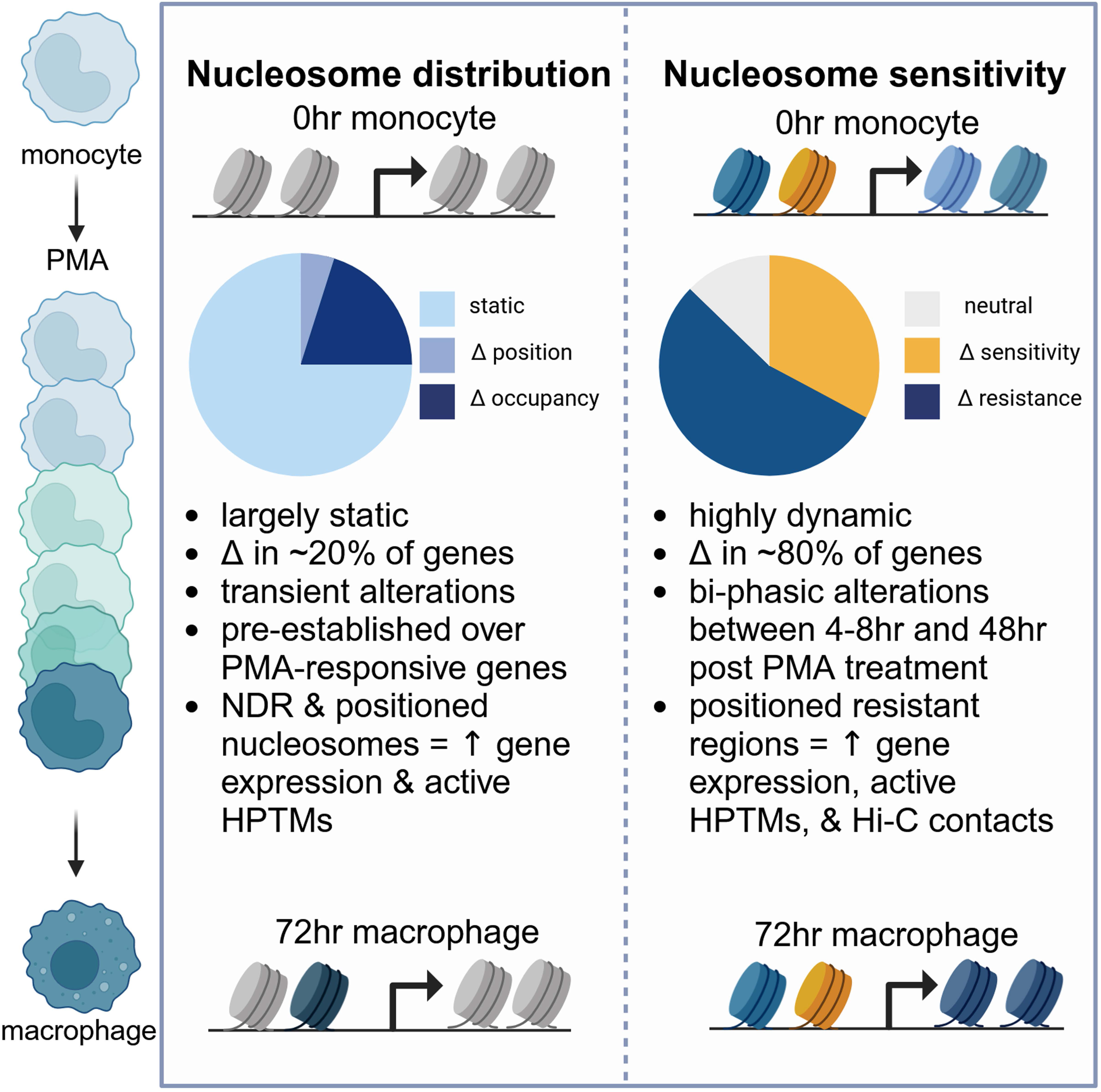
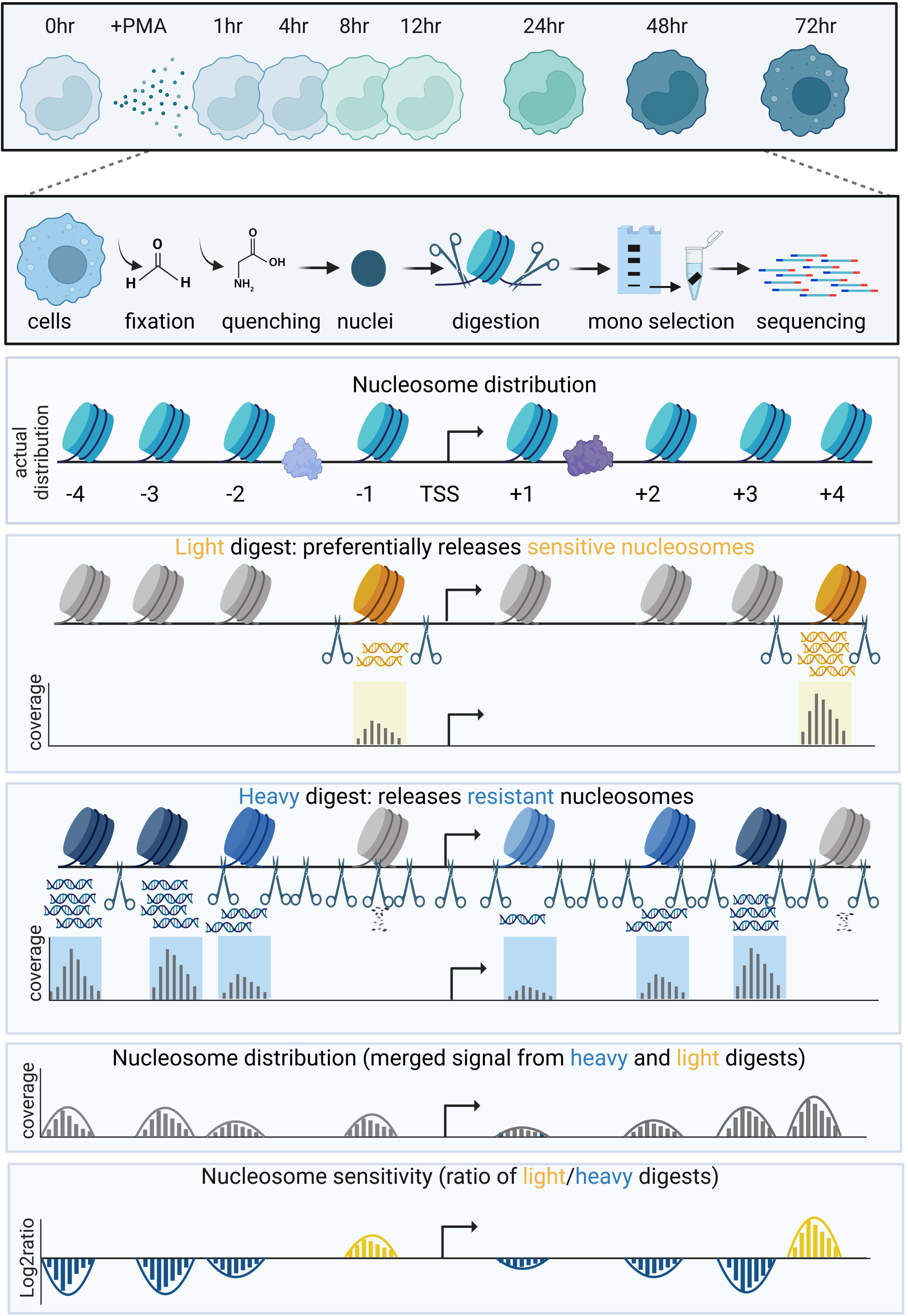
Schematic MNase based assays for measuring nucleosome distribution and sensitivity. THP-1 monocytes were harvested for MNase based chromatin assays at eight timepoints spanning PMA-induced differentiation: 0 hr, 1 hr, 4 hr, 8 hr, 12 hr, 24 hr, 48 hr, and 72 hr. The distribution of nucleosomes within pol II promoters was determined by digesting crosslinked nuclei with two concentrations of MNase. Sensitive nucleosomes were preferentially released under light digest conditions while resistant nucleosomes were preferentially released under heavy digest conditions. Nucleosome distribution was determined by combining nucleosome footprints from both heavy and light digests. The log2 ratio of light/heavy digests was used to determine sensitivity to MNase digestion.

**Figure 2.**
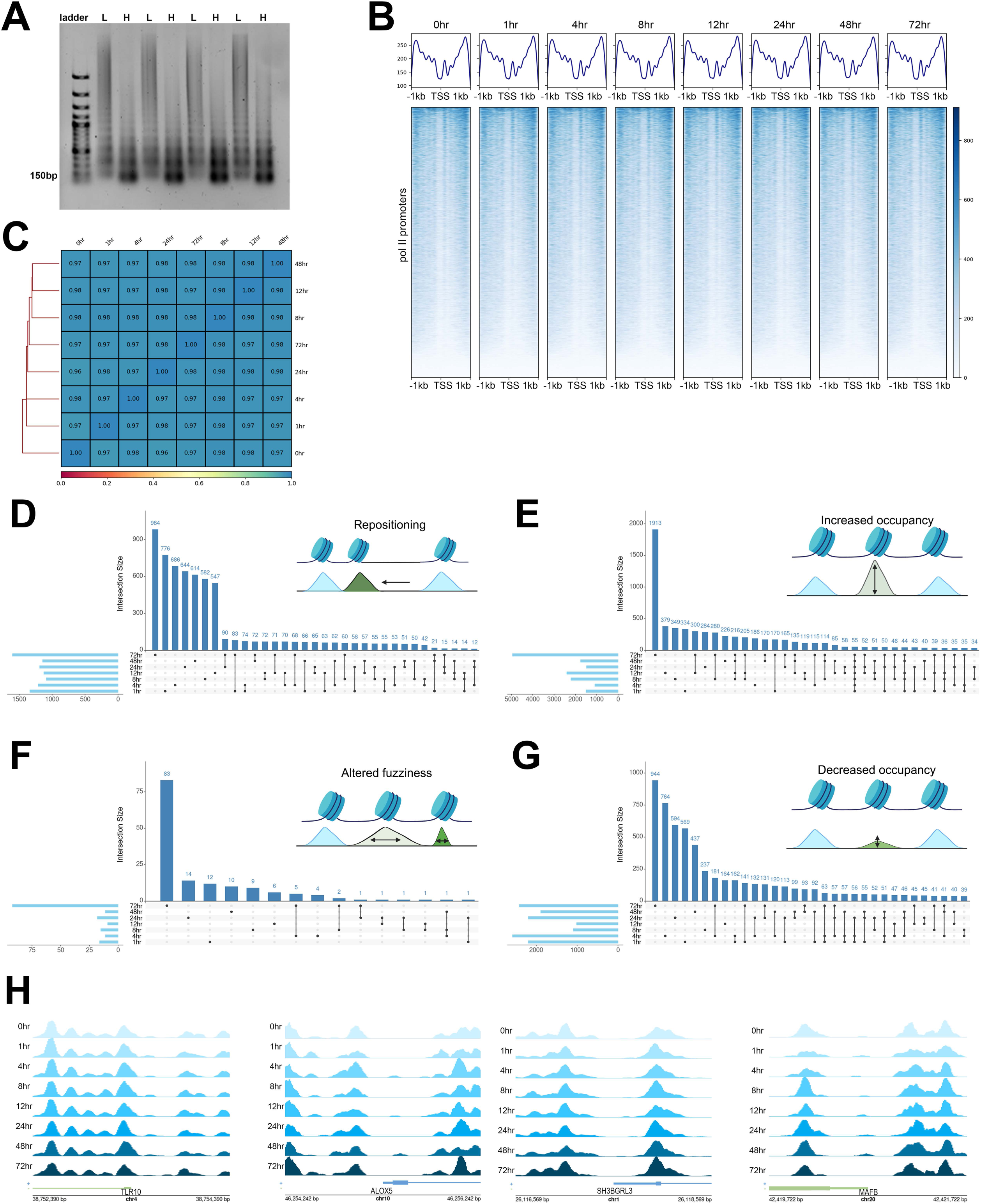
Nucleosome distribution profiles are remarkably stable throughout monocyte differentiation. A. Representative heavy (H) and light (L) digests of crosslinked chromatin run on an agarose gel. B. Heatmaps show nucleosome occupancy signal over a 2kb region centered on the TSS of 21,857 genes. Average signal is shown in plots above each heatmap. Gene order is identical across heatmaps and is determined by the greatest aggregate signal. C. Pearson correlation of nucleosome occupancy in 10bp bins over 2kb promoter regions for all pol II genes. Coefficient values are shown in each square of the heatmap. D. Upset plot showing the relationship between genes with one or more nucleosome with altered position following PMA treatment. The total number of genes for each sample is shown as a pale blue horizontal bar on the left of the plot. Black lines and filled dots connect timepoints with shared genes and the total number of genes in any given combination is reported in dark blue font over the vertical bars. E. Upset plot showing shared and unique genes with altered fuzziness following PMA treatment. F. Upset plot showing shared and unique genes with increased occupancy following PMA treatment. G. Upset plot showing shared and unique genes with decreased occupancy following PMA treatment. H. Nucleosome profiles at all time points over the *TLR10*, *ALOX5, SH3BGRL3,* and *MAFB* promoters.

We used the previously described mTSS-seq promoter capture approach ^42,43^ to enrich RNA pol II promoter sequences for higher resolution nucleosome mapping over 2kb regions flanking the transcription start sites (TSSs) of RefSeq genes. To verify the effectiveness of our approach, we mapped nucleosome occupancy over 4kb regions spanning the targeted promoters (Supp. Fig. 1A). We observed a greater than 100 fold increase in coverage over the targeted 2kb region centered on the TSS compared to the flanking 2kb regions indicating successful enrichment of promoter sequences. This approach produced high-resolution nucleosome maps across pol II promoters with as few as 20 million reads per sample. We next compared nucleosome distribution signal over all promoters at each time point in the PMA-induced differentiation time course. We found that all samples were visually similar with well defined nucleosomes occupying canonical positions flanking a nucleosome depleted region (NDR) in all samples (Fig. 2B). We found that nucleosome distribution was nearly identical across all time points when average signals were directly compared with only minor variations which were not statistically significant (Supp. Fig 1B). Pairwise comparison of nucleosome signal in 10bp bins across 2kb promoter windows genome wide showed Pearson correlation coefficients above 0.96 for all samples (Fig. 2C) further indicating that samples were highly similar across all time points. This high similarity is not unexpected as phenotypically divergent human cell lines also show similar nucleosome distribution profiles (Supp. Fig. 1C, and ^30,42,43,64^) when viewed in aggregate. As these analyses were based on comparison of aggregate promoter signals, we opted for a more comprehensive analysis of nucleosome distribution on a single nucleosome scale. We utilized three measures of nucleosome distribution to identify promoters with alterations in response to PMA induced differentiation.

The first measure was nucleosome repositioning which we defined as any nucleosome with a more than 80bp shift in summit location compared to the undifferentiated sample. We identified between 1087 and 1602 genes with one or more repositioned nucleosomes which met this criterion (Table 1). The majority of these genes were unique to individual time points or common to fewer than three time points suggesting multiple waves of transient repositioning during the monocyte to macrophage transition (Fig. 2D). Gene ontology (GO) analysis using the 2024 Reactome Pathways gene set ^50,51^ revealed a number of known pathways that are implicated in PMA induced differentiation (Supp. Fig. 2). In particular, TRKA activation, calcium signaling, GPCR and ephrin signaling, and lipid metabolism pathways have been associated with THP-1 differentiation and phenotypic alterations in macrophages^36^. Interestingly, the final 72hr time point had the greatest number of repositioned nucleosomes (Table 1), perhaps indicating an additional period of remodeling as cells recover active differentiation pathways and establish a new physiological state.

**Table 1.** Number of genes with one or more re-distributed nucleosomes within 2kb promoter regions following PMA induced differentiation.

| Time | repositioning | altered fuzziness | increased occupancy | decreased occupancy |
| --- | --- | --- | --- | --- |
| 1hr | 1337 | 16 | 1497 | 2194 |
| 4hr | 1209 | 11 | 1084 | 2580 |
| 8hr | 1087 | 15 | 2206 | 1090 |
| 12hr | 1125 | 9 | 2402 | 1011 |
| 24hr | 1189 | 18 | 1476 | 2193 |
| 48hr | 1144 | 11 | 1754 | 1890 |
| 72hr | 1602 | 92 | 4946 | 2413 |

The second measure of nucleosome distribution was nucleosome fuzziness which is related to the broadness or narrowness of a nucleosome footprint. We defined any nucleosome with a greater than 1.5 fold change in fuzziness compared to the 0hr sample combined with an FDR and p-value less than 0.05 as having altered fuzziness. Very few genes had one or more nucleosomes with altered fuzziness at any of the early time points (Table 1). Only the 72hr time point had a substantial number of nucleosomes which met our criteria. Accordingly, it was not unsurprising that the majority of genes with alterations in fuzziness were only observed at single time points rather than shared across time (Fig. 2F). Considering the low gene count, gene ontology was only performed on the 72hr sample which revealed terms including ROS and RNS production, GTPase cycles, Vitamin B1 metabolism, and glycosphingolipid metabolism which are all associated with macrophage differentiation. This suggests that while this measure produced a small number of genes, these genes may have functional relevance in differentiated cells.

The final measure of altered nucleosome distribution was nucleosome occupancy which we classified as increased or decreased occupancy compared to the baseline pattern observed in undifferentiated cells. We defined a change in occupancy as occurring if there was a greater than 5 fold change in occupancy compared to the 0hr sample combined with a p-value and FDR less than 0.05. We observed between 1084 and 4946 genes with one or more nucleosomes within promoter regions with increased occupancy compared to baseline following PMA treatment (Table 1). The majority of these genes were unique to a single time point (Fig. 2E) and the greatest difference was observed in the 72hr sample (Table 1). These genes were enriched for a number of gene ontology terms including differentiation, ERK and JNK pathways, calcium signaling, TP53 regulation, olfactory signaling, and nephron development (Supp. Fig. 4). While some of these pathways (e.g. calcium signaling) have clear relevance to macrophage differentiation, many are apparently unrelated to myeloid development suggesting that widespread alterations in nucleosome occupancy reflect untargeted alterations in many promoters rather than specific remodeling of target genes. We found between 1011 and 2580 genes with one or more nucleosomes within promoter regions with a decrease in nucleosome occupancy following PMA treatment (Table 1). Interestingly, unlike other measures of altered distribution, the 4hr time point had the greatest number of altered genes, not the 72hr sample. The majority of genes with decreased nucleosome occupancy were found at a single time point although there was a trend of many genes common to the 1hr, 4hr and one other time point (Fig. 2G). These genes were enriched for ontology terms including caspase activation, methylation, acyl chain remodeling, ligand-receptor interactions, and olfactory signaling (Supp. Fig. 5).

These measures of nucleosome distribution demonstrate that there are alterations in nucleosome position, fuzziness, and occupancy within a subset of gene promoters and that the greatest alterations are observed in fully differentiated cells. Many of these alterations are transient as they are frequently observed in one or two time points rather than found throughout the differentiation process. These patterns can be observed in single loci examples such as the *TLR10* promoter (Fig. 2H) which represents the near static nucleosome distribution pattern across the differentiation process that was observed over the majority of promoters. In contrast, the *ALOX5* promoter is highly variable with repositioning events at 12hr and 24hr, an occupancy increase at 72hr, and occupancy decreases at 4hr, 12hr, and 48hr (Fig 2H). These alterations coincide with a threefold increase in expression of this potential ferroptosis regulating gene in macrophages^65^. The *SH3BGRL3* promoter has decreased occupancy at the 72hr time point which also coincides with overexpression in macrophages(Fig. 2H). In contrast, the *MAFB* promoter, while visually somewhat variable, did not meet any of our defined cutoffs for nucleosome distribution alterations at any time point. This gene is silent in monocytes and highly expressed in macrophages demonstrating that nucleosome distribution alone is not a sufficient indicator of gene expression. These patterns illustrate the utility of time-resolved analyses to observe transient alterations not captured from before and after comparisons. While we cannot rule out rapid transient alterations in nucleosome distribution occurring between selected timepoints, this result suggests that nucleosome remodeling occurs at specified times on a limited set of promoters during PMA induced monocyte differentiation.

### Nucleosome distribution over PMA-response promoters is pre-established in undifferentiated monocytes

We next examined the relationship between nucleosome distribution and gene expression in undifferentiated monocytes and PMA differentiated macrophages. We organized nucleosome distribution from 0hr and 72hr samples over gene expression quartiles obtained from THP-1 monocytes and PMA-induced fully differentiated macrophages ^14^. We found that the top quartile of highly expressed genes in both monocytes and macrophages was defined by a pronounced nucleosome depleted region (NDR) and highly phased, well positioned flanking nucleosomes (Fig. 3A, quartile 1). In contrast, poorly expressed genes lacked a NDR and had highly occupied nucleosomes across the promoter region (Fig. 3A, quartile 4). These patterns show that distinct nucleosome architectures are present over poorly expressed and highly expressed genes in both monocytes and macrophages.

**Figure 3.**
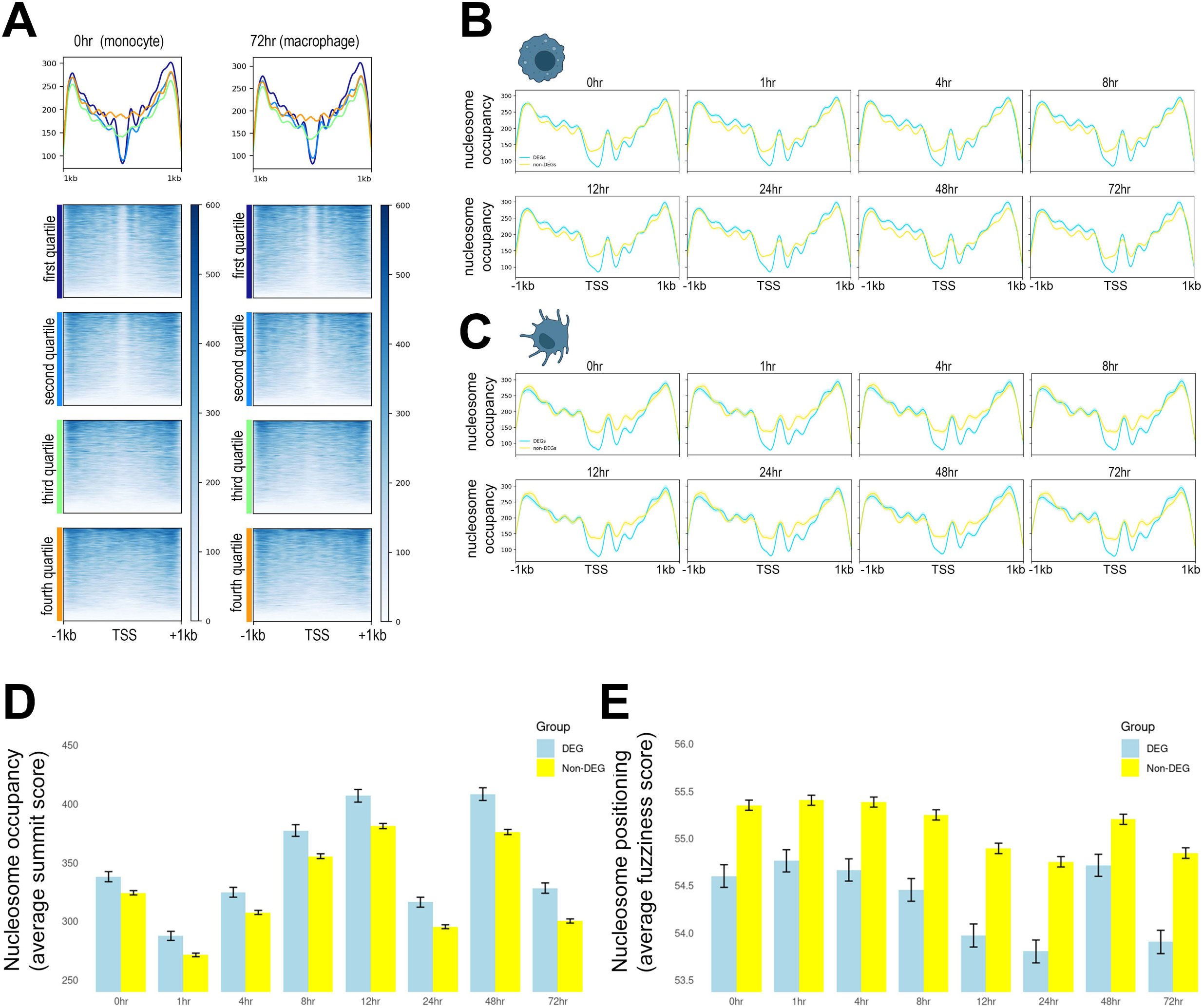
Nucleosome distribution over PMA response gene promoters is pre-established in undifferentiated monocytes. A. Heatmaps show nucleosome distribution over the TSS of pol II genes separated into quartiles based on gene expression in undifferentiated monocytes (0hr sample) and PMA differentiated macrophages (72hr sample). B. Average THP-1 nucleosome distribution plotted over PMA-responsive DEGs and a matched set of non-DEG promoters at all time points following PMA treatment. 95% confidence intervals are shown in shaded regions behind average lines. C. Average THP-1 nucleosome distribution plotted over monocyte-derived dendritic cells DEGs and a matched set of non-DEG promoters during the monocyte to macrophage transition. 95% confidence intervals are shown in shaded regions behind the average lines. D. Average nucleosome summit scores for all nucleosomes within PMA-responsive DEG and non-DEG promoters with 95% confidence intervals for all time points. F. Average nucleosome fuzziness scores with 95% confidence intervals for all nucleosomes within PMA-responsive DEG and non-DEG promoters for all time points.

We next investigated whether PMA-responsive genes exhibit any unique features in the monocytes, prior to and during their differentiation. For this, we examined the promoter states of 3,209 genes defined as differentially expressed genes (DEGs) in the PMA-induced differentiation of monocytes to macrophages. Promoter profiles were highly consistent over these genes over the differentiation time course and they consistently exhibited a more distinct NDR and a more defined +1 nucleosome, as illustrated by the more pronounced amplitude and definition of the +1 peak, compared to a matched set of non-DEGs (Fig. 3B). These results suggests that the nucleosome promoter architecture, in terms of both positioning and occupancy, of macrophage specific genes is preestablished in monocytes prior to the addition of the differentiation stimulus. We tested this idea by plotting nucleosome distribution over the promoters of DEGs from monocyte derived dendritic cells which are a distinct immune cell type also derived from the myeloid lineage. The dendritic cell DEGs promoters were similarly marked by a more distinct nucleosome depleted region and a strongly positioned +1 nucleosome at all timepoints (Fig. 3C). This suggests that a subset of lineage specific promoters are poised in the undifferentiated monocyte for rapid expression when stimulated by the appropriate cell fate signals. Thus, the nucleosome distribution profile is a reflection of the future potential of a cell rather than the current state alone. To quantify these observations, we used DANPOS3 to calculate both nucleosome summit scores, reflecting occupancy, and fuzziness scores, reflecting positioning, for DEGs and a matched set of non-DEGs. We found that nucleosomes in DEG promoters exhibited significantly higher summit scores, reflecting higher occupancy, and lower fuzziness scores, reflecting stronger phasing, than non-DEGs a (Fig. 3D-E). These trends were observed in monocytes, fully differentiated macrophages, and across all intermediate time points further indicating that this architecture is established in monocytes and maintained across the differentiation process.

### Nucleosome sensitivity to MNase digestion is highly dynamic during differentiation and offers insight into genome potential

Given that measurements of nucleosome distribution were remarkably consistent at the majority of promoters across timepoints, we turned to an analysis of chromatin sensitivity through the differentiation process. We used the log2 ratio of light to heavy digests to define promoter regions which were sensitive and resistant to MNase digestion. Regions which were more abundant in light digests are defined as sensitive to MNase while regions which are overrepresented in heavy digests are defined as resistant to MNase (summarized in Fig. 1).

We mapped sensitivity over the promoters of PMA-treated monocytes at the same time points previously described to determine if susceptibility to MNase would be an informative measure of differentiation-specific chromatin dynamics. We used the untreated THP-1 monocytes (0hr sample) as a basal comparator for changes in nucleosome sensitivity throughout the monocyte to macrophage transition. We utilized kmeans clustering to identify groups of promoters based on dominant sensitivity features in undifferentiated cells (Fig. 4A, 0hr) and traced the sensitivity profiles of these same genes at all other time points (Fig. 4A, 4hr-72hr). This enabled us to identify three distinct promoter classes in undifferentiated cells.

**Figure 4.**
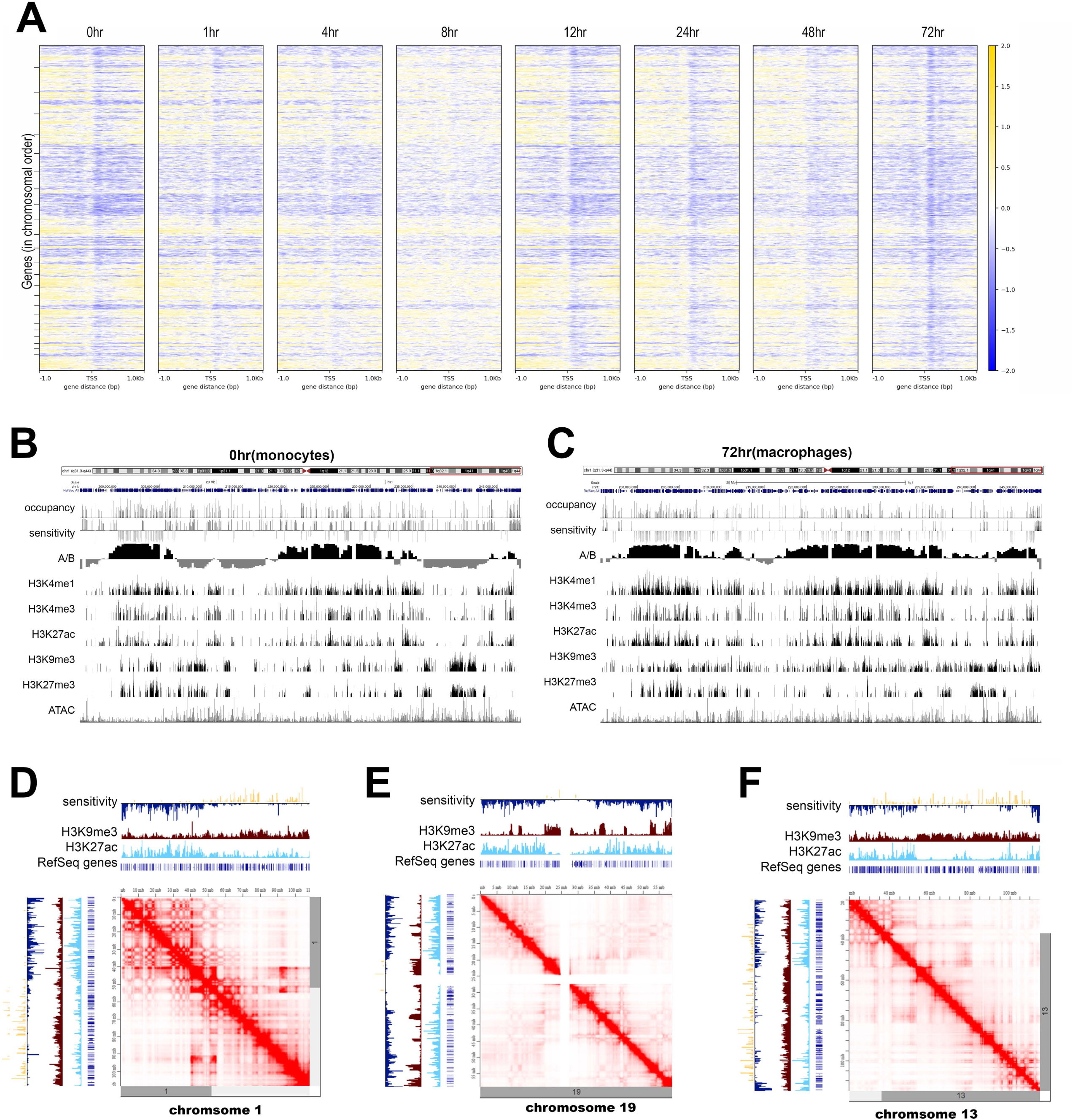
The monocyte to macrophage transition is marked by a dynamic MNase sensitivity and an increase in resistance to MNase digestion. A. Heatmaps show MNase sensitivity over a 2kb region centered on the TSS of pol II promoters during the monocyte to macrophage transition. Promoters are separated into dominant classes by kmeans clustering of the 0hr time point with sensitive regions in gold and resistant regions in blue. Line plots showing average signal for each cluster are shown above heatmaps. Clusters are color-coded and labeled along the Y axis B. Independently clustered heatmaps of sensitivity signal over a 2kb window centered on the TSS for all time points. Gene order and promoter clusters were independently generated for each time point. C. Nucleosome sensitivity profiles over the *IL5, LTB,* and *NFKB1* promoters.

First, a cluster of promoters defined by a diffusely sensitive pattern across the entire 2kb region with no other discernible features (Fig. 4A, 0hr cluster 1). A second promoter class contained a mixture of both weakly sensitive and weakly resistant regions with a mean signal value close to zero (Fig. 4A, 0hr cluster 2). The final promoter class was defined by a strongly positioned resistant nucleosome which occupied regions associated with canonical positioned -2, -1, +1, +2, or +3 positions (Fig. 4A, 0hr clusters 3-7). These results, consistent with our previous work ^30,64^ suggest two main characteristics of human gene promoters: (1) that sensitive nucleosomes do not generally take on discrete positions at gene promoters, and (2) that resistant, but not sensitive, nucleosomes are found at discrete canonical positions in gene promoters.

Next, we wanted to investigate how these patterns of delocalized sensitivity and discretely positioned resistant nucleosomes defined in the untreated THP-1 cells may change over the monocyte-to-macrophage transition. The same gene order defined by the 0hr clusters were applied to all other time points (Fig. 4A, 1hr-72hr) to show alterations in sensitivity over time. There was an immediate disruption of the basal resistant promoter classes within 1hr of PMA induced differentiation (Fig. 4A, 1hr clusters 4-7). This disruption was also apparent at subsequent time points (Fig. 4A, compare clusters 4, 5, 6, and 7 in 0hr to adjacent 1hr, 4hr and 8hr timepoints). This is followed by a transient but partial return to the undifferentiated pattern at 12-72 hours. Promoters in the diffusely sensitive class were more stable over time with similar patterns observed at all timepoints with a subtle decrease in signal between 4-8 hours followed by a recovery in later time points (Fig. 4A, cluster 1 all time points). The mixed promoter class was also largely stable over time with an appreciable increase in resistance over the +1 nucleosome in later time points (Fig. 4A, cluster 2 all time points). The data suggests that this global remodeling is occurring in a biphasic manner, with 0-12 hours appearing as a first phase and a second between 24-72 hours.

The observable disruption of both sensitivity and resistance patterns with a trend towards increasing resistance in macrophages was supported by a statistical analysis of sensitive and resistant nucleosomes across promoters. The number of nucleosomes with differential summit scores in light and heavy digests, as defined by a log fold change greater than 1.5 and FDR and p-values under 0.05 in a DANPOS3 comparison, were compared over time (Table 2). Any nucleosome not meeting these criteria was defined as neutral. By this analysis, 12.4% of nucleosomes were identified as sensitive in undifferentiated monocytes. This number decreased to 10.9% at 8hr following PMA treatment and partially recovered to 11.0% in fully differentiated macrophages. In contrast, 12.5% of nucleosomes were classified as resistant in undifferentiated monocytes. This number dropped rapidly after PMA treatment with only 6.2% of nucleosomes identified as resistant at 8hr post PMA treatment. This proportion increased to 14.4% of promoter nucleosomes in fully differentiated cells 72hr after PMA treatment (Table 2). These results are further supported by a correlation analysis which showed weak correlation values across samples and identified the 8hr and 72hr timepoints as outliers (Supp. Fig. 6A).

**Table 2.** Number and percent of sensitive and resistant nucleosomes within promoters at each time point following PMA induced differentiation. Nucleosomes with a log2 fold change in summit score greater than 1.5 combined with a p-value and FDR under 0.05 were classified as sensitive if more abundant in light digests and resistant if more abundant in heavy digests. All nucleosomes not meeting these thresholds were classified as neutral.

| time | # sensitive | # resistant | # neutral | % sensitive | % resistant |
| --- | --- | --- | --- | --- | --- |
| 0hr | 20018 | 20082 | 121041 | 12.4 | 12.5 |
| 1hr | 18704 | 16824 | 132326 | 11.1 | 10.0 |
| 4hr | 16323 | 12312 | 113796 | 11.5 | 8.6 |
| 8hr | 16164 | 9273 | 123154 | 10.9 | 6.2 |
| 12hr | 20113 | 16344 | 108668 | 13.9 | 11.3 |
| 24hr | 17617 | 15510 | 115879 | 11.8 | 10.4 |
| 48hr | 17947 | 11246 | 121367 | 11.9 | 7.5 |
| 72hr | 16857 | 22058 | 114661 | 11.0 | 14.4 |

To determine the full range of promoter sensitivity dynamics throughout the monocyte to macrophage transition, we next independently generated kmeans clustering maps of nucleosome sensitivity at each timepoint (Fig. 4B). This analysis enabled identification of a new class of promoter that was not observed in undifferentiated monocytes. A class of promoters with a single well positioned and highly sensitive nucleosome just upstream of the immediate TSS was observed in 1hr, 4hr, 8hr, and 24hr samples (Fig. 4B, cluster 2, selected time points). An intermediate type promoter class with a broadly sensitive upstream region was observed at 12hr and 24hr (Fig. 4B, cluster 2 12hr & 48hr). Neither of these classes were present in monocytes (0hr) or fully differentiated macrophages (72hr). Multiple positioned resistant promoter classes were identified at all time points and were consistent with positioning observed in monocytes (Fig. 4B, clusters 3-7). Analysis of the promoters within each cluster during the differentiation time course showed that the diffusely sensitive (Supp. Fig. 6B) and positioned resistant clusters (Supp. Fig. 6D) shared many common genes throughout differentiation while the positioned sensitive (Supp. Fig. 6C) and mixed (Supp. Fig. 6E) clusters were primarily composed of unique genes across different time points. This result emphasizes the highly dynamic nature of nucleosome sensitivity and suggests that common regulatory mechanisms may apply to conserved gene clusters which are not disrupted by these alterations.

To further illustrate these dynamics, we examined specific loci characteristic of the highly dynamic nature of nucleosome sensitivity throughout the differentiation process. The TSS proximal region of the *IL5* promoter is initially highly sensitive upstream of the TSS, becomes less sensitive between 4-8hr after PMA stimulation, and is weakly sensitive in fully differentiated macrophages (Fig. 4F). Similarly time-dependent chromatin dynamics is seen over the *LTB* and *NFKB1* promoters (Fig. 4F). The *LTB* promoter transitions from largely resistant in monocytes to more sensitive during differentiation and back to resistant in macrophages. The *NFKB1* proximal promoter is highly sensitive until 24hr after PMA stimulation and adopts a new resistant architecture in fully differentiated macrophages. Similar dynamics can be observed across promoters with a general trend of disrupted sensitivity between 4hr-8hr and 12hr-48hr post PMA induced differentiation with the establishment of a new sensitivity pattern in fully differentiated macrophages at 72hr.

### Promoters with a single discretely positioned resistant nucleosome are associated with active chromatin marks and increased gene expression

Considering the widespread nature of sensitivity dynamics during differentiation, we were interested in the relationship between sensitivity and gene expression. We compared nucleosome sensitivity profiles at promoters of genes separated into expression quartiles. We found that highly expressed genes showed a district sensitivity pattern with a sensitive region just upstream of the immediate TSS flanked by highly resistant regions in both monocytes and macrophages (Supp. Fig. 7A, quartiles 1-2). This contrasts with poorly expressed genes which lack defined sensitive and resistant regions across the promoter (Supp. Fig. 7B, quartiles 3-4) in both monocytes and macrophages. These trends were also observed over PMA-response genes which are overexpressed in PMA-derived macrophages (Supp. Fig. 7B). Response genes were sensitive just upstream of the TSS and resistant over the +1 nucleosome. This pattern was disrupted during the 4-8hr and 48hr periods previously described, and was most pronounced 72hr after PMA stimulation (Supp. Fig. 7B). Interestingly, plotting nucleosome sensitivity over dendritic cell DEGs shows a similar pattern (Supp. Fig. 7C) demonstrating the widespread nature of sensitivity alterations which impact the majority of promoters. These findings show that, during differentiation, nucleosome sensitivity dynamics are pronounced, biphasic, and not isolated to macrophage specific genes. In contrast to nucleosome distribution, a basal nucleosome sensitivity organization does not appear to be preestablished in the undifferentiated monocyte.

Given that highly expressed genes were associated with characteristic MNase sensitivity dynamics, we were interested in the relationship between sensitivity and other dynamic genomic features including accessibility and histone modifications. In order to further understand the interplay between chromatin sensitivity and other well described characteristics of promoter architecture, we compared MNase sensitivity with nucleosome distribution, genome accessibility (ATAC-seq), and histone modifications, specifically H3K4me3 and H3K27ac, which are known markers of active promoters in undifferentiated monocytes (Fig. 5A) and differentiated macrophages (Fig. 5B). Promoters were organized into dominant classes defined by sensitivity patterns at each time point. When compared to maps of nucleosome distribution, promoters containing dominantly positioned resistant regions are marked by well defined nucleosome depleted regions over the TSS flanked by strongly positioned -1 and +1 nucleosomes (Fig. 5A-B, clusters 3-7). In contrast, the highly sensitive promoter class showed highly occupied but poorly positioned nucleosomes lacking a defined nucleosome depleted region (Fig. 5A-B, cluster 1). Maps of promoter accessibility as measured by ATAC-seq were not clearly correlated with measurements of sensitivity to MNase in undifferentiated monocytes as ATAC signal was nearly uniform over all promoter classes (Fig. 5A). ATAC-seq signal was somewhat enriched in positioned resistant clusters in fully differentiated macrophages (Fig. 5B, greatest signal in clusters 3-6 based on average plots) This observation demonstrates that ATAC-seq and MNase-sensitivity are measuring different and distinct aspects of promoter organization. Active histone marks, H3K4me3 and H3K27ac, were highly enriched in promoters with well-positioned resistant nucleosomes, suggesting that promoters containing these discrete resistant regions are active or poised promoters in both monocytes and macrophages (Fig. 5A, clusters 3-7; Fig. 5B, clusters 2-7; most clearly seen in average plots above heatmaps). Consistent with these observations,the clusters containing discrete resistant locations also showed elevated gene expression in monocytes (Fig. 5C) and macrophages (Fig. 5D). This work shows that a positioned resistant nucleosome is a dynamic feature of promoter architecture. Furthermore, this positioned resistant nucleosome is an important epigenomic characteristic of active promoters. Interestingly, a cluster of diffusely sensitive promoters in monocytes (Fig. 5A, cluster 1) was associated with weakly positioned nucleosomes, low H3K4me3 and H3K27ac coverage, and the lowest gene expression of all the clusters. These diffusely sensitive promoters were also observed as part of cluster 1 in differentiated macrophages (Fig. 5B) where they were similarly associated with low HPTM coverage and reduced expression (Fig. 5D). This result suggests that diffuse sensitivity to enzymatic digestion can be used to define inactive promoters.

**Figure 5.**
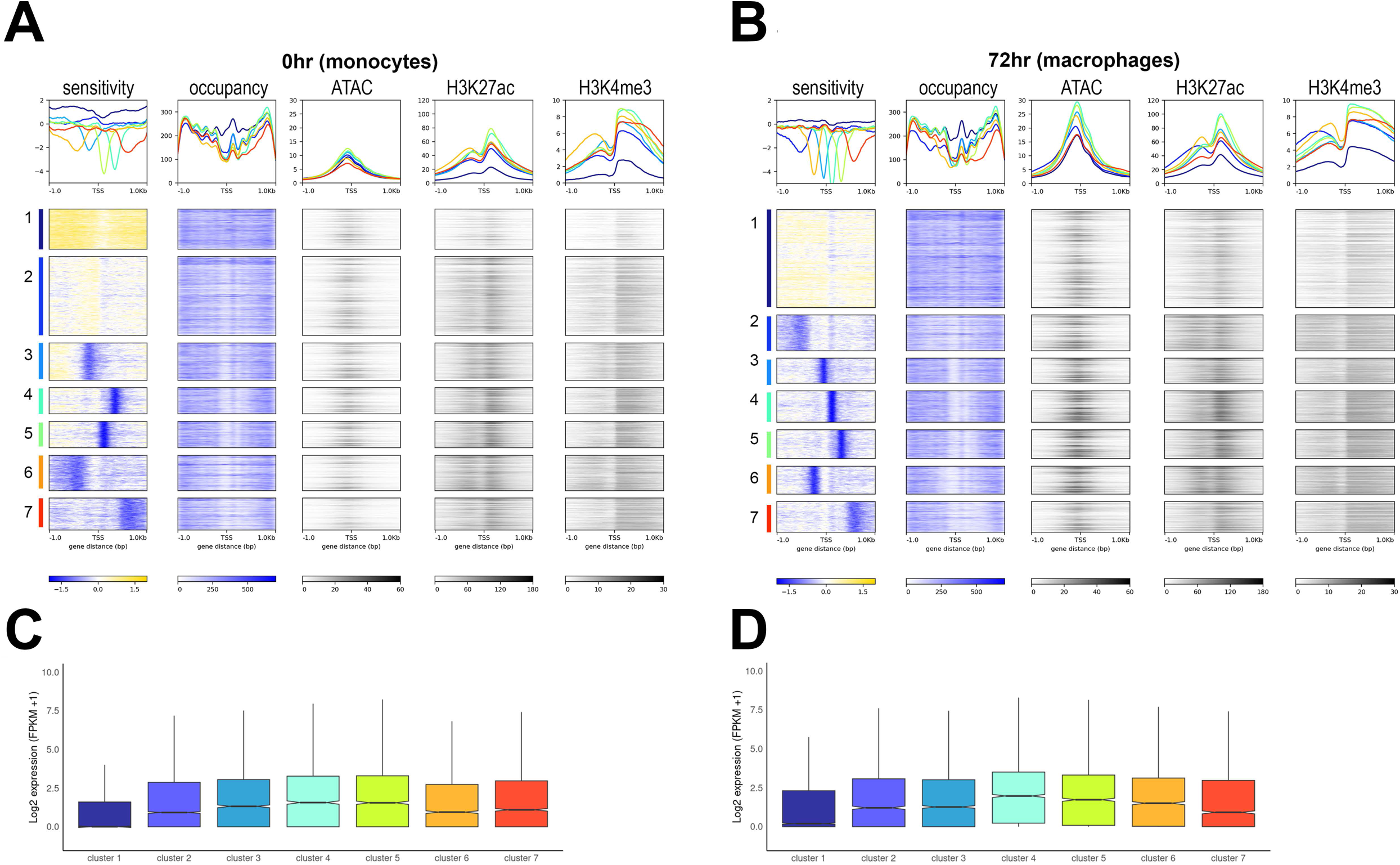
Positioned resistant nucleosomes are associated with active chromatin marks and increased gene expression in both monocytes and macrophages. A. Heatmaps of MNase sensitivity, nucleosome occupancy, ATACseq, H3K27ac, and H3K4me3 ChIP signal over pol II promoters in monocytes (0hr) grouped by sensitivity clusters. Gene order is identical across all maps in the series. Average plots are shown above each heatmap with color coding based on clusters listed along the y-axis. B. Heatmaps of MNase sensitivity, nucleosome occupancy, ATACseq, H3K27ac, and H3K4me3 ChIP signal over pol II promoters in macrophages (72hr) grouped by sensitivity clusters. Gene order is identical across all maps in the series. Average plots are shown above each heatmap with color coding based on clusters listed along the y-axis. C. Boxplots show gene expression (log2 (FPKM+1)) for each monocyte sensitivity cluster shown in 5A. Notches indicate the median expression value and the middle 50% of the data displayed as the box, and the whiskers extend to the smallest and largest values within 1.5 times the IQR. D. Boxplots show gene expression (log2 (FPKM+1)) for each macrophage sensitivity cluster shown in 5B.

Monocytes have a mixed cluster that is characterized by weak sensitive and resistant signals (Fig. 5A cluster 2), which is an intermediate between the active clusters characterized by positioned resistant nucleosomes (Fig. 5A clusters 3-7) and inactive clusters characterized by delocalized sensitive nucleosomes (Fig. 5A cluster 1). The fully differentiated macrophage cells do not generate a distinct mixed cluster (Fig. 5B) despite numerous clustering attempts. This suggests that terminal differentiation may involve the resolution of intermediate chromatin states into more distinct resistant or sensitive promoter architectures, contributing to stable chromatin profiles in the differentiated cell. This transition from an intermediate to a resistant state can be seen in the substantial increase from 9,345 promoters containing a positioned resistant nucleosome in monocytes to 13,539 promoters in macrophages. These results demonstrate that nucleosome sensitivity to nuclease provides new defining features of active and inactive chromatin at the resolution of individual nucleosomes at promoters undergoing PMA induced monocyte differentiation.

### Nucleosome sensitivity occurs in domain like patterns on a chromosomal scale and is associated with higher order chromatin features

The classification and dynamics of MNase sensitivity of individual promoters during the monocyte-to-macrophage transition led us to ask if sensitivity to MNase could characterize megabase scale domains of the chromosomes or was limited primarily to the individual nucleosome scale. To answer this, we plotted promoter sensitivity across all promoters in chromosomal order (Fig. 6A). This analysis identified domains of promoters exhibiting sensitivity or resistance to MNase on a kilobase to megabase scale as indicated by distinct patterns of highly sensitive or highly resistant promoters within individual chromosomes. These sensitivity patterns showed dynamic alterations after PMA induced differentiation with an initial disruption between 4-8hr and a trend towards increasing resistance in fully differentiated cells (Fig. 6A, compare sharpness of patterns in 0hr vs 8hr vs 72hr). Closer analysis of chromosome 19 showed that while nucleosome distribution patterns (Supp. Fig. 8A) were highly stable throughout differentiation on a chromosomal scale, nucleosome sensitivity patterns were more variable and trended towards increased resistance in macrophages (Supp. Fig. 8B). Interestingly, sensitivity patterns co-localized with both Hi-C compartments and HPTM domains in both monocytes and macrophages (Fig. 6B-C) in genome browser views of individual chromosomes. Predominantly resistant regions overlapped with the A compartment and active HPTMs while more sensitive regions corresponded with the inactive B compartment and repressive HPTMS (Fig. 6B-C). The colocalization between resistant regions and active HPTMs was also observed when plotting heatmaps in chromosomal order in both unstimulated monocytes and differentiated macrophages (Supp. Fig. 8C-D).

**Figure 6.**
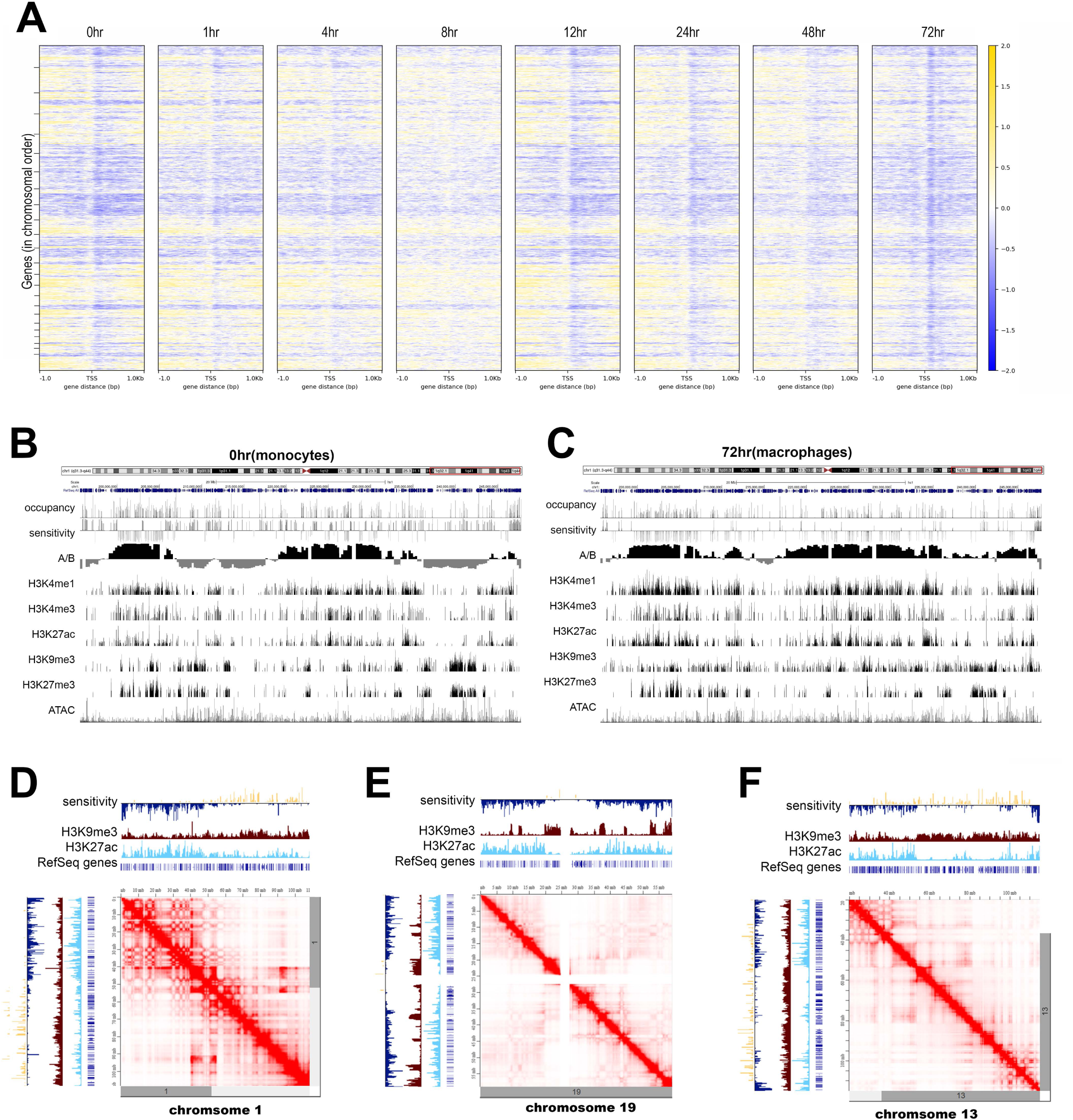
Differentiation induced nucleosome sensitivity dynamics track with known epigenomic features. A.Heatmaps of nucleosome sensitivity over pol II promoters in chromosomal order from chr1-chrY for all time points. Tick marks indicate approximate chromosome sizes. Gene order is identical for all maps in the series. B. Genome browser view of nucleosome occupancy, sensitivity, Hi-C A/B compartments, H3K4me1, H3K4me3, H3K27ac, H3K9me3, H3K27me3, and ATACseq over a 40mB region of chromosome 1 in unstimulated monocytes (0hr). C. Genome browser view of nucleosome occupancy, sensitivity, Hi-C A/B compartments, H3K4me1, H3K4me3, H3K27ac, H3K9me3, H3K27me3, and ATACseq over a 40mB region of chromosome 1 in macrophages (72hr). D. Overlay of sensitivity (yellow/blue), H3K9me3 (burgundy), and H3K27ac (cyan) trucks over a Hi-C map (red) of a 110mb window of chromosome 1 in unstimulated monocytes (0hr). E. Overlay of sensitivity (yellow/blue), H3K9me3 (burgundy), and H3K27ac (cyan) trucks over a Hi-C map (red) of the entire chromosome 19 in unstimulated monocytes (0hr). F. Overlay of sensitivity (yellow/blue), H3K9me3 (burgundy), and H3K27ac (cyan) trucks over a Hi-C map (red) of a 100mb window of chromosome 13 in unstimulated monocytes (0hr).

Given the observed domain-like structure of these MNase sensitivity patterns across chromosomes, we hypothesized that sensitivity might correspond to A/B compartments identified in Hi-C experiments. To test this, we compared MNase sensitivity maps to Hi-C data from THP-1 monocytes and macrophages ^14^. We found that nucleosome sensitivity corresponds to Hi-C A/B compartments (Fig. 6B-C) and increased 3D contacts (Fig. 6D-F). Regions which are highly resistant fall within the active Hi-C A compartment and tend to have increased contacts and more active HPTMs. In contrast, regions which are highly sensitive correspond with the inactive B compartment, have fewer intrachromosomal contacts, and increased heterochromatin marks. These findings suggest that nucleosome sensitivity measures features linked to higher-order chromatin structure, with resistant regions corresponding to more transcriptionally active A compartments, and sensitive regions aligning with inactive B compartments and heterochromatin. This reinforces the idea that nucleosome sensitivity is influenced by, and contributes to, the overall chromosomal architecture.

## Discussion

This study presents the first high resolution temporal analysis of promoter nucleosome architecture dynamics during the PMA induced monocyte differentiation. Previous studies have highlighted extensive changes in epigenetic marks and chromatin structure during this transition ^5,14,58,66,67^, but they primarily focused on a direct comparison between monocytes and macrophages without examining intermediate stages in the differentiation process. Our findings reveal that, while nucleosome distribution remains stable over more than 80% of promoters, nucleosome sensitivity to MNase digestion is highly dynamic throughout differentiation. These sensitivity patterns correlate with active chromatin marks, suggesting that nucleosome sensitivity, combined with targeted nucleosome remodeling, plays a key role in regulating gene expression during differentiation. Moreover, our study highlights the interplay between promoter sensitivity and higher-order chromatin structure. The techniques described here are broadly applicable and offer valuable insights into chromatin architecture in a wide range of tissues and cell types.

Our analysis shows that promoter nucleosome positioning remains largely unchanged across the monocyte-to-macrophage differentiation process (Fig. 2A). Despite substantial transcriptomic and phenotypic changes, nucleosome occupancy and phasing at most promoters exhibit minimal variation. We observed transient changes in nucleosome distribution at selected promoters, suggesting localized nucleosome remodeling at key regulatory loci as cells differentiate. Notably, we found that promoters of PMA-responsive differentially expressed genes (DEGs) exhibited increased nucleosome occupancy and stronger positioning even before differentiation (Fig. 3B). This pre-established landscape in monocytes suggests that macrophage-specific promoter architectures are primed for rapid gene activation when the appropriate differentiation stimulus is detected, a finding that underscores the role of chromatin in cell-type-specific gene regulation. Actively expressed promoters consistently showed stronger nucleosome positioning, clearer phasing, and well-defined nucleosome-depleted regions over the TSS, distinguishing them from poorly expressed or silent genes in both monocytes and macrophages.

In contrast to the near static nature of nucleosome positioning, nucleosome sensitivity exhibited striking temporal variation. Notably, the global remodeling of sensitivity at key time points (4-8 hr and 48 hr) after PMA induced differentiation suggest that these may represent critical windows in the differentiation process (Fig. 4A). These findings suggest that nucleosome positioning serves as a stable scaffold, while sensitivity represents a dynamic mechanism allowing or restricting access to underlying genomic regions as differentiation progresses. The observed biphasic sensitivity pattern may reflect different regulatory stages of macrophage differentiation: an early phase that initiates the broader differentiation program, followed by a later phase that stabilizes the differentiated state. This sensitivity remodeling affected the majority of promoters, implying a genome-wide phenomenon that extends beyond highly expressed promoters. These insights underscore the central role of nucleosome sensitivity in regulating access to the genome during differentiation.

Beyond nucleosome positioning, our study underscores the role of nucleosome sensitivity to MNase as a crucial marker of promoter activity. Highly expressed genes were associated with well-positioned, resistant nucleosomes and active histone marks such as H3K4me3 and H3K27ac, while diffusely sensitive nucleosomes define inactive promoters (Fig. 5A-B) in both monocytes and macrophages. This work supports our previous observation of this association between resistance and active promoters in near normal epithelial cells ^30^. This adds a new layer of understanding to promoter regulation, emphasizing the functional significance of chromatin sensitivity in gene expression.

Interestingly, differentiated macrophages lacked the mixed sensitive/resistant nucleosome patterns seen in monocytes, suggesting a progression toward more defined chromatin states during differentiation. Moreover, domain-wide patterns of nucleosome sensitivity across chromosomes mirrored Hi-C A/B compartments, suggesting a connection between local promoter dynamics and higher-order chromatin organization (Fig. 6). This result is particularly interesting since our measure of MNase sensitivity is based solely on a 2kb region at the promoters of protein-coding genes while Hi-C utilizes data from the entire genome. These results highlight the complexity of chromatin architecture during differentiation and suggest that nucleosome sensitivity, along with stable nucleosome positioning, plays a pivotal role in modulating gene activity and chromatin organization on a genome-wide scale.

Our investigation demonstrates that nucleosome distribution and sensitivity can be a revealing measure of functional chromatin and has provided valuable insights into the nuclear landscape. Our combination of these measurements with a time-resolved study has enabled the identification of critical chromatin transitions in the differentiation process and offers a detailed view of the regulatory steps governing these transitions. While our study focuses on the ordered events characterizing the monocyte to macrophage transition, it is noteworthy that the approach holds promise in characterizing cellular transitions across diverse biological phenomena, ranging from differentiation processes to implications in cancer and responses to various stimuli. We expect that further studies of this nature will enable a more holistic understanding of chromatin dynamics and its regulatory role in cellular processes.

## ACKNOWLEDGEMENTS

We thank Lorea Arambarri, Shahin Behrouz Sharif, and Mahdi Khadem for constructive conversations regarding data analysis and all members of the Dennis and Bass labs for critiquing figures and results.

## AUTHOR CONTRIBUTIONS

JMB: Conceptualization, Methodology, Formal Analysis, Visualization, Software, Data curation, Writing--original draft and editing. BDB: Conceptualization, Methodology, Writing—review. MK: Writing—review. HWB: Methodology, Writing—review and editing. JHD: Funding acquisition, Conceptualization, Supervision, Resources, Writing – review & editing.

## DATA AVAILABILITY

All raw and processed sequencing data generated in this study have been submitted to the NCBI Gene Expression Omnibus (GEO; https://www.ncbi.nlm.nih.gov/geo/) under accession number GSE254274.

