## Supplementary material for "Transient alterations in nucleosome distribution and sensitivity to nuclease define the THP-1 monocyte to macrophage transition": Supp_figs

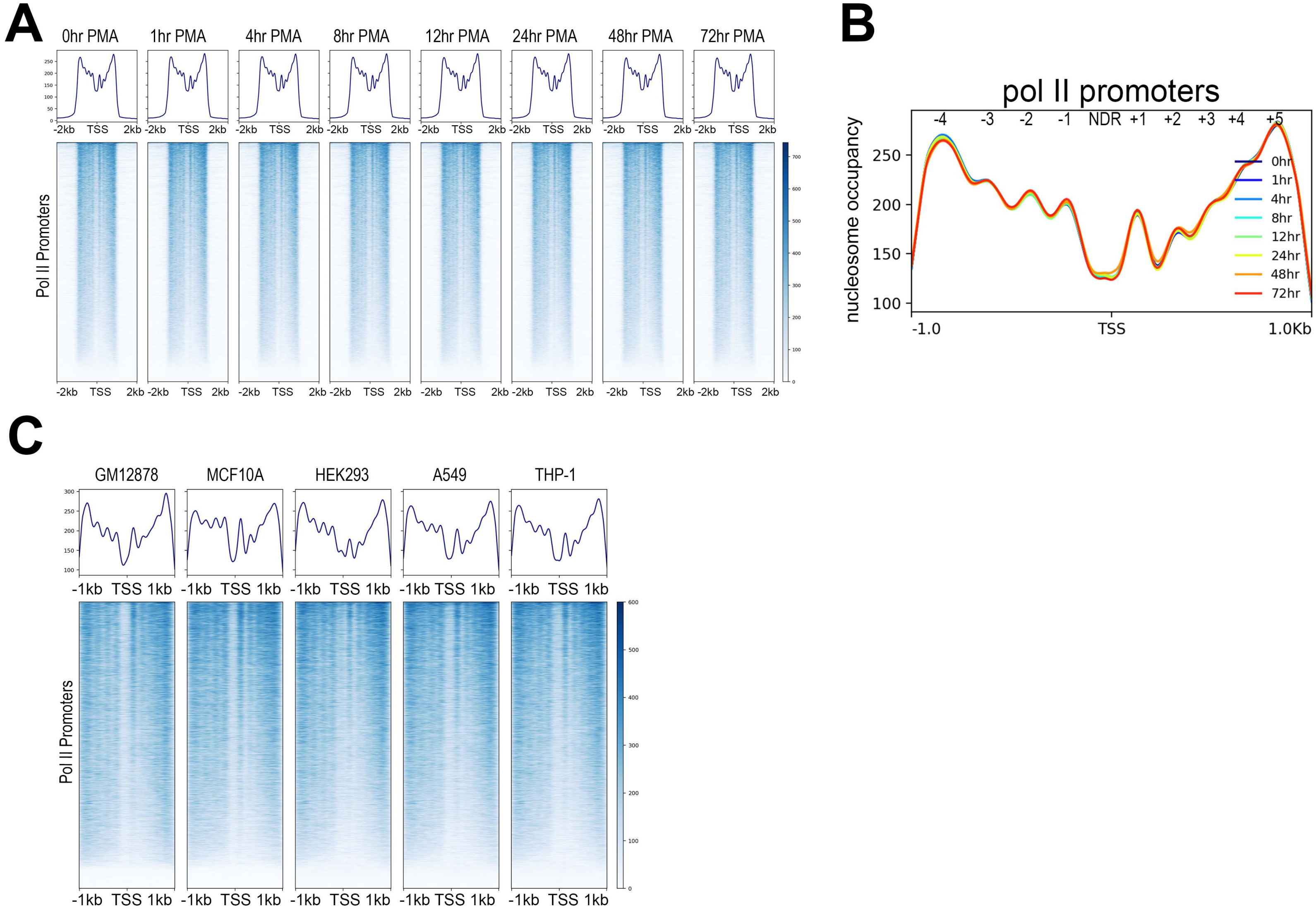
A
0hr PMA
1hr PMA
4hr PMA
8hr PMA
12hr PMA
24hr PMA
48hr PMA
72hr PMA
-2kb TSS 2kb
Pol II Promoters
B
pol II promoters
-4 -3 -2 -1 NDR +1 +2 +3 +4 +5
nucleosome occupancy
0hr
1hr
4hr
8hr
12hr
24hr
48hr
72hr
-1.0
TSS
1.0Kb
C
GM12878
MCF10A
HEK293
A549
THP-1
-1kb TSS 1kb
Pol II Promoters

### Supp Fig 1. Efficiency of promoter capture and nucleosome distribution profiles across treatments and cell types

A. Heatmaps showing nucleosome occupancy signal over a 4kb window centered on the TSS of 21, 857 pol II genes. Averaged coverage profiles above each heatmap show enrichment of reads within 2kb of the TSS. B. Average nucleosome occupancy overlayed at all timepoints with 95% confidence intervals shown in pale shading. Nucleosome positions and NDR are labeled above each feature. C. Heatmaps showing nucleosome occupancy over 2kb centered on the TSS of pol II genes in GM12878, MCF10A, HEK293, A549, and undifferentiated THP-1 cells. Average plots are shown above each heatmap. Gene order is identical in all samples and determined by maximum signal values across all cell lines.

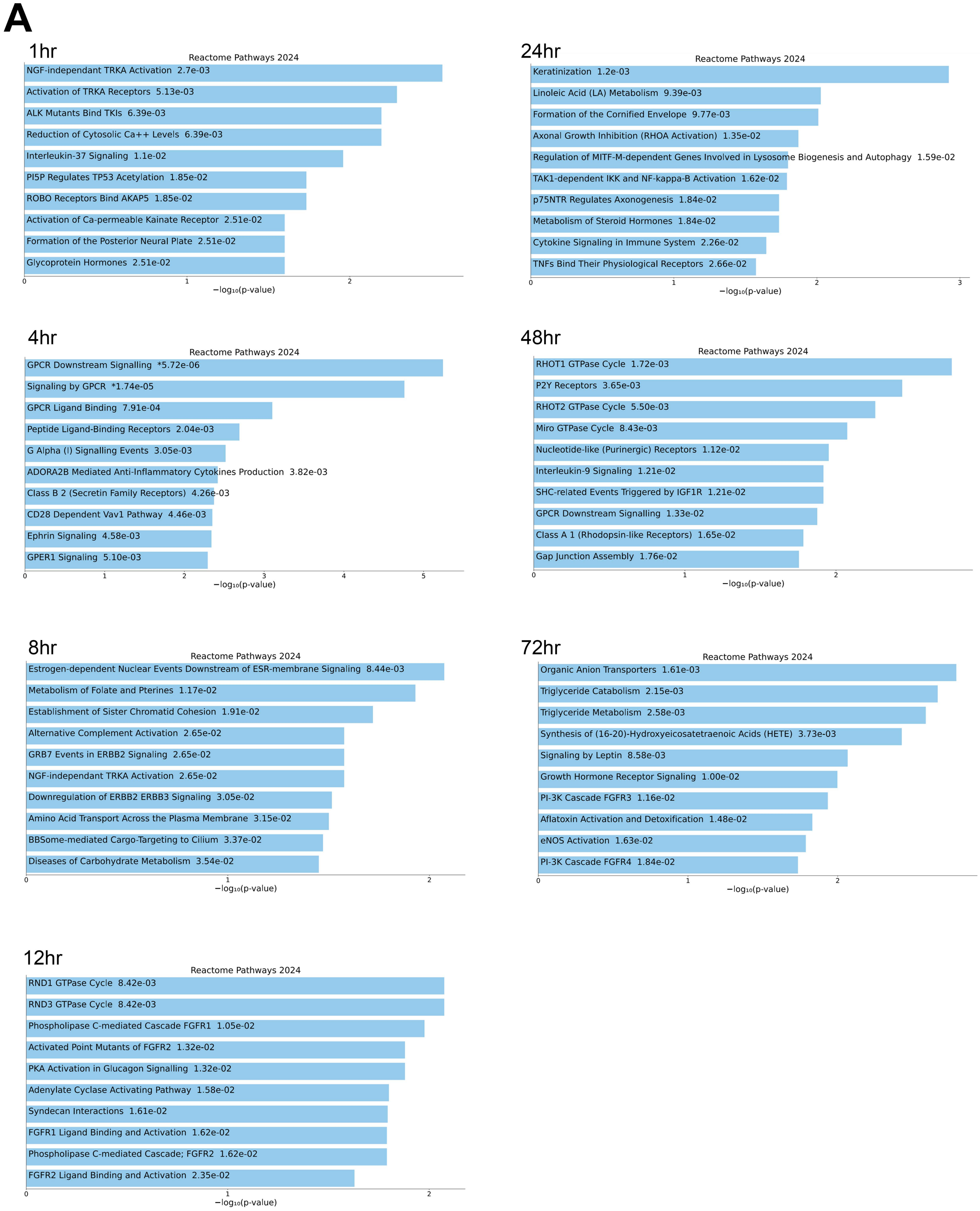
A
1hr
Reactome Pathways 2024
NGF-independant TRKA Activation 2.7e-03
Activation of TRKA Receptors 5.13e-03
ALK Mutants Bind TKIs 6.39e-03
Reduction of Cytosolic Ca++ Levels 6.39e-03
Interleukin-37 Signaling 1.1e-02
PI5P Regulates TP53 Acetylation 1.85e-02
ROBO Receptors Bind AKAP5 1.85e-02
Activation of Ca-permeable Kainate Receptor 2.51e-02
Formation of the Posterior Neural Plate 2.51e-02
Glycoprotein Hormones 2.51e-02
0
1
2
−log₁₀(p-value)
24hr
Reactome Pathways 2024
Keratinization 1.2e-03
Linoleic Acid (LA) Metabolism 9.39e-03
Formation of the Cornified Envelope 9.77e-03
Axonal Growth Inhibition (RHOA Activation) 1.35e-02
Regulation of MITF-M-dependent Genes Involved in Lysosome Biogenesis and Autophagy 1.59e-02
TAK1-dependent IKK and NF-kappa-B Activation 1.62e-02
p75NTR Regulates Axonogenesis 1.84e-02
Metabolism of Steroid Hormones 1.84e-02
Cytokine Signaling in Immune System 2.26e-02
TNFs Bind Their Physiological Receptors 2.66e-02
0
1
2
3
−log₁₀(p-value)
4hr
Reactome Pathways 2024
GPCR Downstream Signalling *5.72e-06
Signaling by GPCR *1.74e-05
GPCR Ligand Binding 7.91e-04
Peptide Ligand-Binding Receptors 2.04e-03
G Alpha (I) Signalling Events 3.05e-03
ADORA2B Mediated Anti-Inflammatory Cytokines Production 3.82e-03
Class B 2 (Secretin Family Receptors) 4.26e-03
CD28 Dependent Vav1 Pathway 4.46e-03
Ephrin Signaling 4.58e-03
GPER1 Signaling 5.10e-03
0
1
2
3
4
5
−log₁₀(p-value)
48hr
Reactome Pathways 2024
RHOT1 GTPase Cycle 1.72e-03
P2Y Receptors 3.65e-03
RHOT2 GTPase Cycle 5.50e-03
Miro GTPase Cycle 8.43e-03
Nucleotide-like (Purinergic) Receptors 1.12e-02
Interleukin-9 Signaling 1.21e-02
SHC-related Events Triggered by IGF1R 1.21e-02
GPCR Downstream Signalling 1.33e-02
Class A 1 (Rhodopsin-like Receptors) 1.65e-02
Gap Junction Assembly 1.76e-02
0
1
2
−log₁₀(p-value)
8hr
Reactome Pathways 2024
Estrogen-dependent Nuclear Events Downstream of ESR-membrane Signaling 8.44e-03
Metabolism of Folate and Pterines 1.17e-02
Establishment of Sister Chromatid Cohesion 1.91e-02
Alternative Complement Activation 2.65e-02
GRB7 Events in ERBB2 Signaling 2.65e-02
NGF-independant TRKA Activation 2.65e-02
Downregulation of ERBB2 ERBB3 Signaling 3.05e-02
Amino Acid Transport Across the Plasma Membrane 3.15e-02
BBSome-mediated Cargo-Targeting to Cilium 3.37e-02
Diseases of Carbohydrate Metabolism 3.54e-02
0
1
2
−log₁₀(p-value)
72hr
Reactome Pathways 2024
Organic Anion Transporters 1.61e-03
Triglyceride Catabolism 2.15e-03
Triglyceride Metabolism 2.58e-03
Synthesis of (16-20)-Hydroxyeicosatetraenoic Acids (HETE) 3.73e-03
Signaling by Leptin 8.58e-03
Growth Hormone Receptor Signaling 1.00e-02
PI-3K Cascade FGFR3 1.16e-02
Aflatoxin Activation and Detoxification 1.48e-02
eNOS Activation 1.63e-02
PI-3K Cascade FGFR4 1.84e-02
0
1
2
−log₁₀(p-value)
12hr
Reactome Pathways 2024
RND1 GTPase Cycle 8.42e-03
RND3 GTPase Cycle 8.42e-03
Phospholipase C-mediated Cascade FGFR1 1.05e-02
Activated Point Mutants of FGFR2 1.32e-02
PKA Activation in Glucagon Signalling 1.32e-02
Adenylate Cyclase Activating Pathway 1.58e-02
Syndecan Interactions 1.61e-02
FGFR1 Ligand Binding and Activation 1.62e-02
Phospholipase C-mediated Cascade; FGFR2 1.62e-02
FGFR2 Ligand Binding and Activation 2.35e-02
0
1
2
−log₁₀(p-value)

### Supp. Fig. 2. Gene ontology for promoters with altered positioning after PMA treatment

A. Top GO terms for genes with altered nucleosome positioning at each time point following PMA treatment. Adjusted P values are shown beside ontology terms. Bars in blue indicate significant enrichments.

A

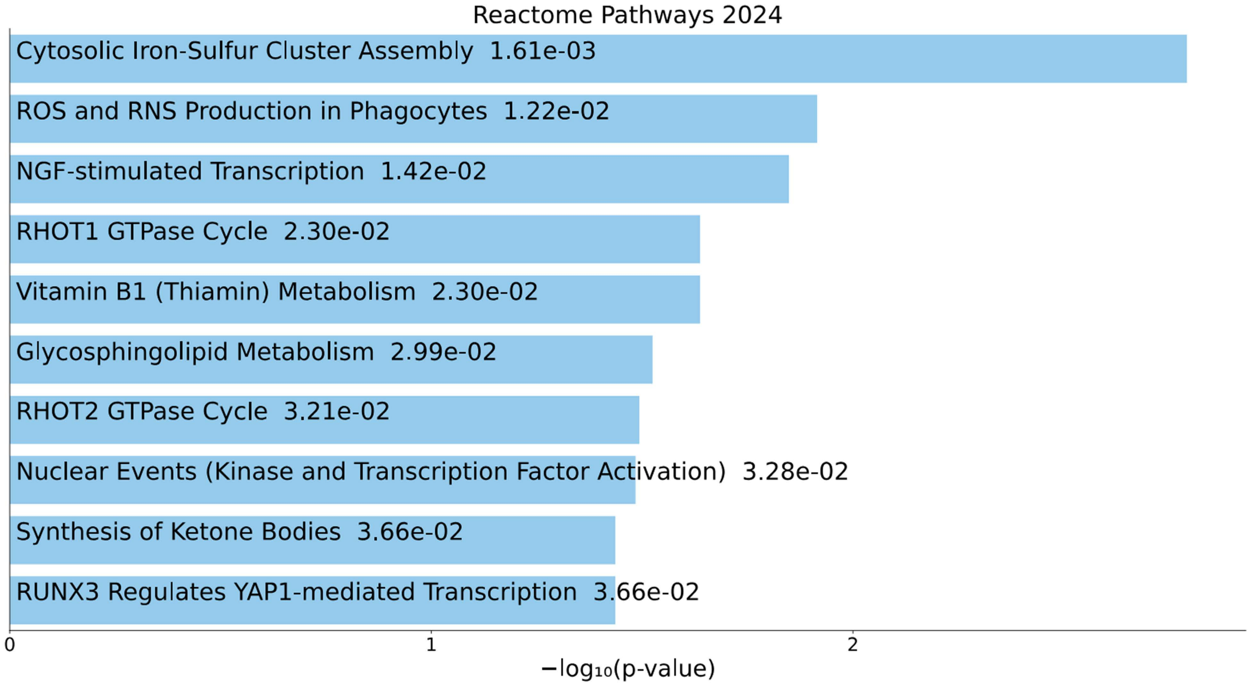
Reactome Pathways 2024
Cytosolic Iron-Sulfur Cluster Assembly 1.61e-03
ROS and RNS Production in Phagocytes 1.22e-02
NGF-stimulated Transcription 1.42e-02
RHOT1 GTPase Cycle 2.30e-02
Vitamin B1 (Thiamin) Metabolism 2.30e-02
Glycosphingolipid Metabolism 2.99e-02
RHOT2 GTPase Cycle 3.21e-02
Nuclear Events (Kinase and Transcription Factor Activation) 3.28e-02
Synthesis of Ketone Bodies 3.66e-02
RUNX3 Regulates YAP1-mediated Transcription 3.66e-02
0
1
2
−log10(p-value)

### Supp. Fig. 3. Gene ontology for promoters with altered fuzziness after PMA treatment

A. Top GO terms for genes with altered nucleosome fuzziness at each time point following PMA treatment. Adjusted P values are shown beside ontology terms. Bars in blue indicate significant enrichments.

A

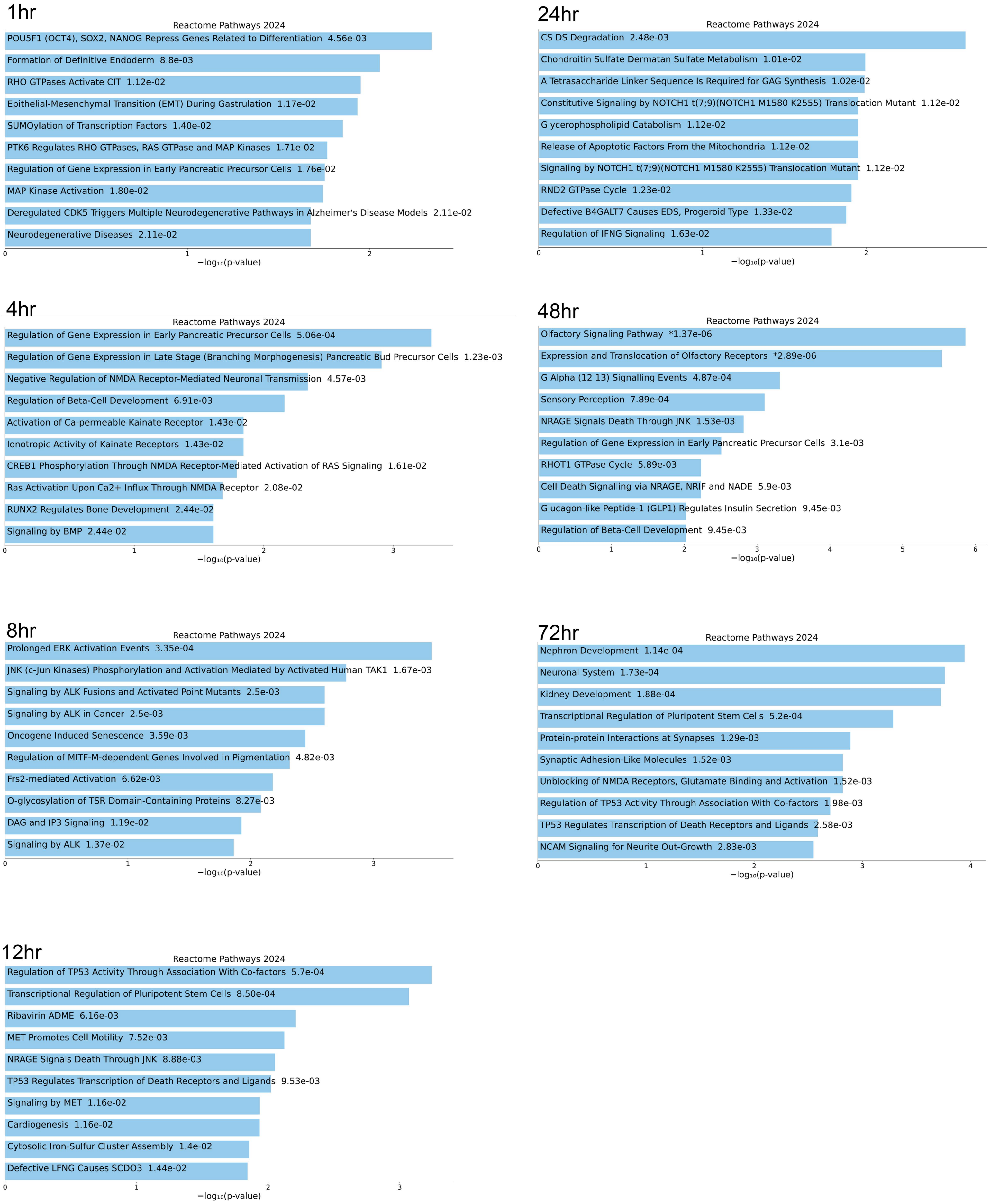
1hr
Reactome Pathways 2024
POU5F1 (OCT4), SOX2, NANOG Repress Genes Related to Differentiation 4.56e-03
Formation of Definitive Endoderm 8.8e-03
RHO GTPases Activate CIT 1.12e-02
Epithelial-Mesenchymal Transition (EMT) During Gastrulation 1.17e-02
SUMOylation of Transcription Factors 1.40e-02
PTK6 Regulates RHO GTPases, RAS GTPase and MAP Kinases 1.71e-02
Regulation of Gene Expression in Early Pancreatic Precursor Cells 1.76e-02
MAP Kinase Activation 1.80e-02
Deregulated CDK5 Triggers Multiple Neurodegenerative Pathways in Alzheimer's Disease Models 2.11e-02
Neurodegenerative Diseases 2.11e-02
0
1
2
−log₁₀(p-value)
24hr
Reactome Pathways 2024
CS DS Degradation 2.48e-03
Chondroitin Sulfate Dermatan Sulfate Metabolism 1.01e-02
A Tetrasaccharide Linker Sequence Is Required for GAG Synthesis 1.02e-02
Constitutive Signaling by NOTCH1 t(7;9)(NOTCH1 M1580 K2555) Translocation Mutant 1.12e-02
Glycerophospholipid Catabolism 1.12e-02
Release of Apoptotic Factors From the Mitochondria 1.12e-02
Signaling by NOTCH1 t(7;9)(NOTCH1 M1580 K2555) Translocation Mutant 1.12e-02
RND2 GTPase Cycle 1.23e-02
Defective B4GALT7 Causes EDS, Progeroid Type 1.33e-02
Regulation of IFNG Signaling 1.63e-02
0
1
2
−log₁₀(p-value)
4hr
Reactome Pathways 2024
Regulation of Gene Expression in Early Pancreatic Precursor Cells 5.06e-04
Regulation of Gene Expression in Late Stage (Branching Morphogenesis) Pancreatic Bud Precursor Cells 1.23e-03
Negative Regulation of NMDA Receptor-Mediated Neuronal Transmission 4.57e-03
Regulation of Beta-Cell Development 6.91e-03
Activation of Ca-permeable Kainate Receptor 1.43e-02
Ionotropic Activity of Kainate Receptors 1.43e-02
CREB1 Phosphorylation Through NMDA Receptor-Mediated Activation of RAS Signaling 1.61e-02
Ras Activation Upon Ca2+ Influx Through NMDA Receptor 2.08e-02
RUNX2 Regulates Bone Development 2.44e-02
Signaling by BMP 2.44e-02
0
1
2
3
−log₁₀(p-value)
48hr
Reactome Pathways 2024
Olfactory Signaling Pathway *1.37e-06
Expression and Translocation of Olfactory Receptors *2.89e-06
G Alpha (12 13) Signalling Events 4.87e-04
Sensory Perception 7.89e-04
NRAGE Signals Death Through JNK 1.53e-03
Regulation of Gene Expression in Early Pancreatic Precursor Cells 3.1e-03
RHOT1 GTPase Cycle 5.89e-03
Cell Death Signalling via NRAGE, NRIF and NADE 5.9e-03
Glucagon-like Peptide-1 (GLP1) Regulates Insulin Secretion 9.45e-03
Regulation of Beta-Cell Development 9.45e-03
0
1
2
3
4
5
6
−log₁₀(p-value)
8hr
Reactome Pathways 2024
Prolonged ERK Activation Events 3.35e-04
JNK (c-Jun Kinases) Phosphorylation and Activation Mediated by Activated Human TAK1 1.67e-03
Signaling by ALK Fusions and Activated Point Mutants 2.5e-03
Signaling by ALK in Cancer 2.5e-03
Oncogene Induced Senescence 3.59e-03
Regulation of MITF-M-dependent Genes Involved in Pigmentation 4.82e-03
Frs2-mediated Activation 6.62e-03
O-glycosylation of TSR Domain-Containing Proteins 8.27e-03
DAG and IP3 Signaling 1.19e-02
Signaling by ALK 1.37e-02
0
1
2
3
−log₁₀(p-value)
72hr
Reactome Pathways 2024
Nephron Development 1.14e-04
Neuronal System 1.73e-04
Kidney Development 1.88e-04
Transcriptional Regulation of Pluripotent Stem Cells 5.2e-04
Protein-protein Interactions at Synapses 1.29e-03
Synaptic Adhesion-Like Molecules 1.52e-03
Unblocking of NMDA Receptors, Glutamate Binding and Activation 1.52e-03
Regulation of TP53 Activity Through Association With Co-factors 1.98e-03
TP53 Regulates Transcription of Death Receptors and Ligands 2.58e-03
NCAM Signaling for Neurite Out-Growth 2.83e-03
0
1
2
3
4
−log₁₀(p-value)
12hr
Reactome Pathways 2024
Regulation of TP53 Activity Through Association With Co-factors 5.7e-04
Transcriptional Regulation of Pluripotent Stem Cells 8.50e-04
Ribavirin ADME 6.16e-03
MET Promotes Cell Motility 7.52e-03
NRAGE Signals Death Through JNK 8.88e-03
TP53 Regulates Transcription of Death Receptors and Ligands 9.53e-03
Signaling by MET 1.16e-02
Cardiogenesis 1.16e-02
Cytosolic Iron-Sulfur Cluster Assembly 1.4e-02
Defective LFNG Causes SCDO3 1.44e-02
0
1
2
3
−log₁₀(p-value)

### Supp. Fig. 4. Gene ontology for promoters with increased occupancy after PMA treatment

A.  Top GO terms for genes with increased nucleosome occupancy at each time point following PMA treatment. Adjusted P values are shown beside ontology terms. Bars in blue indicate significant enrichments.

A

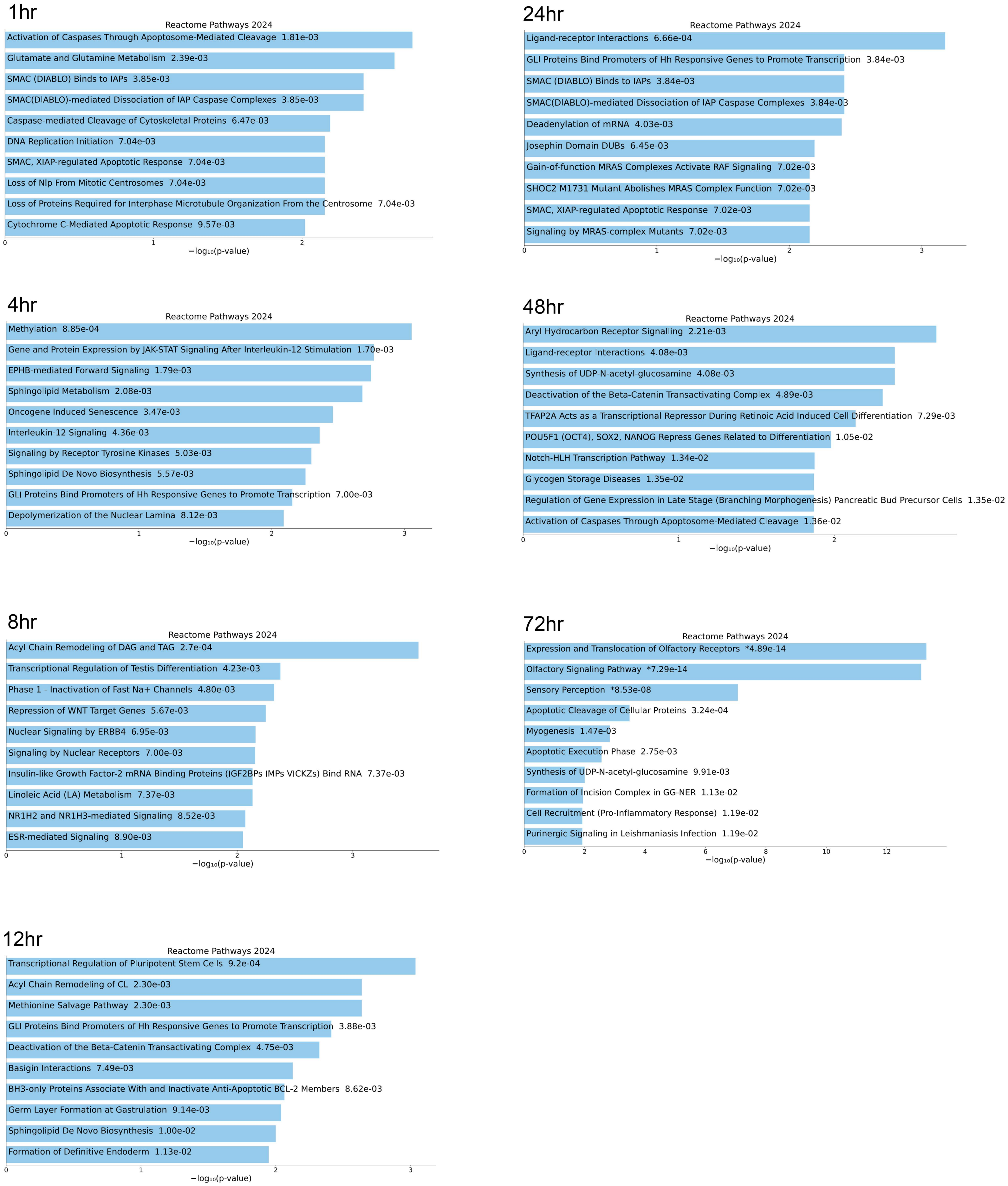
1hr
Reactome Pathways 2024
Activation of Caspases Through Apoptosome-Mediated Cleavage 1.81e-03
Glutamate and Glutamine Metabolism 2.39e-03
SMAC (DIABLO) Binds to IAPs 3.85e-03
SMAC(DIABLO)-mediated Dissociation of IAP Caspase Complexes 3.85e-03
Caspase-mediated Cleavage of Cytoskeletal Proteins 6.47e-03
DNA Replication Initiation 7.04e-03
SMAC, XIAP-regulated Apoptotic Response 7.04e-03
Loss of Nlp From Mitotic Centrosomes 7.04e-03
Loss of Proteins Required for Interphase Microtubule Organization From the Centrosome 7.04e-03
Cytochrome C-Mediated Apoptotic Response 9.57e-03
0
1
2
−log₁₀(p-value)
24hr
Reactome Pathways 2024
Ligand-receptor Interactions 6.66e-04
GLI Proteins Bind Promoters of Hh Responsive Genes to Promote Transcription 3.84e-03
SMAC (DIABLO) Binds to IAPs 3.84e-03
SMAC(DIABLO)-mediated Dissociation of IAP Caspase Complexes 3.84e-03
Deadenylation of mRNA 4.03e-03
Josephin Domain DUBs 6.45e-03
Gain-of-function MRAS Complexes Activate RAF Signaling 7.02e-03
SHOC2 M1731 Mutant Abolishes MRAS Complex Function 7.02e-03
SMAC, XIAP-regulated Apoptotic Response 7.02e-03
Signaling by MRAS-complex Mutants 7.02e-03
0
1
2
3
−log₁₀(p-value)
4hr
Reactome Pathways 2024
Methylation 8.85e-04
Gene and Protein Expression by JAK-STAT Signaling After Interleukin-12 Stimulation 1.70e-03
EPHB-mediated Forward Signaling 1.79e-03
Sphingolipid Metabolism 2.08e-03
Oncogene Induced Senescence 3.47e-03
Interleukin-12 Signaling 4.36e-03
Signaling by Receptor Tyrosine Kinases 5.03e-03
Sphingolipid De Novo Biosynthesis 5.57e-03
GLI Proteins Bind Promoters of Hh Responsive Genes to Promote Transcription 7.00e-03
Depolymerization of the Nuclear Lamina 8.12e-03
0
1
2
3
−log₁₀(p-value)
48hr
Reactome Pathways 2024
Aryl Hydrocarbon Receptor Signalling 2.21e-03
Ligand-receptor Interactions 4.08e-03
Synthesis of UDP-N-acetyl-glucosamine 4.08e-03
Deactivation of the Beta-Catenin Transactivating Complex 4.89e-03
TFAP2A Acts as a Transcriptional Repressor During Retinoic Acid Induced Cell Differentiation 7.29e-03
POU5F1 (OCT4), SOX2, NANOG Repress Genes Related to Differentiation 1.05e-02
Notch-HLH Transcription Pathway 1.34e-02
Glycogen Storage Diseases 1.35e-02
Regulation of Gene Expression in Late Stage (Branching Morphogenesis) Pancreatic Bud Precursor Cells 1.35e-02
Activation of Caspases Through Apoptosome-Mediated Cleavage 1.36e-02
0
1
2
−log₁₀(p-value)
8hr
Reactome Pathways 2024
Acyl Chain Remodeling of DAG and TAG 2.7e-04
Transcriptional Regulation of Testis Differentiation 4.23e-03
Phase 1 - Inactivation of Fast Na+ Channels 4.80e-03
Repression of WNT Target Genes 5.67e-03
Nuclear Signaling by ERBB4 6.95e-03
Signaling by Nuclear Receptors 7.00e-03
Insulin-like Growth Factor-2 mRNA Binding Proteins (IGF2BPs IMPs VICKZs) Bind RNA 7.37e-03
Linoleic Acid (LA) Metabolism 7.37e-03
NR1H2 and NR1H3-mediated Signaling 8.52e-03
ESR-mediated Signaling 8.90e-03
0
1
2
3
−log₁₀(p-value)
72hr
Reactome Pathways 2024
Expression and Translocation of Olfactory Receptors *4.89e-14
Olfactory Signaling Pathway *7.29e-14
Sensory Perception *8.53e-08
Apoptotic Cleavage of Cellular Proteins 3.24e-04
Myogenesis 1.47e-03
Apoptotic Execution Phase 2.75e-03
Synthesis of UDP-N-acetyl-glucosamine 9.91e-03
Formation of Incision Complex in GG-NER 1.13e-02
Cell Recruitment (Pro-Inflammatory Response) 1.19e-02
Purinergic Signaling in Leishmaniasis Infection 1.19e-02
0
2
4
6
8
10
12
−log₁₀(p-value)
12hr
Reactome Pathways 2024
Transcriptional Regulation of Pluripotent Stem Cells 9.2e-04
Acyl Chain Remodeling of CL 2.30e-03
Methionine Salvage Pathway 2.30e-03
GLI Proteins Bind Promoters of Hh Responsive Genes to Promote Transcription 3.88e-03
Deactivation of the Beta-Catenin Transactivating Complex 4.75e-03
Basigin Interactions 7.49e-03
BH3-only Proteins Associate With and Inactivate Anti-Apoptotic BCL-2 Members 8.62e-03
Germ Layer Formation at Gastrulation 9.14e-03
Sphingolipid De Novo Biosynthesis 1.00e-02
Formation of Definitive Endoderm 1.13e-02
0
1
2
3
−log₁₀(p-value)

### Supp. Fig. 5. Gene ontology for promoters with decreased occupancy after PMA treatment

A. Top GO terms for genes with decreased nucleosome occupancy at each time point following PMA treatment. Adjusted P values are shown beside ontology terms. Bars in blue indicate significant enrichments.

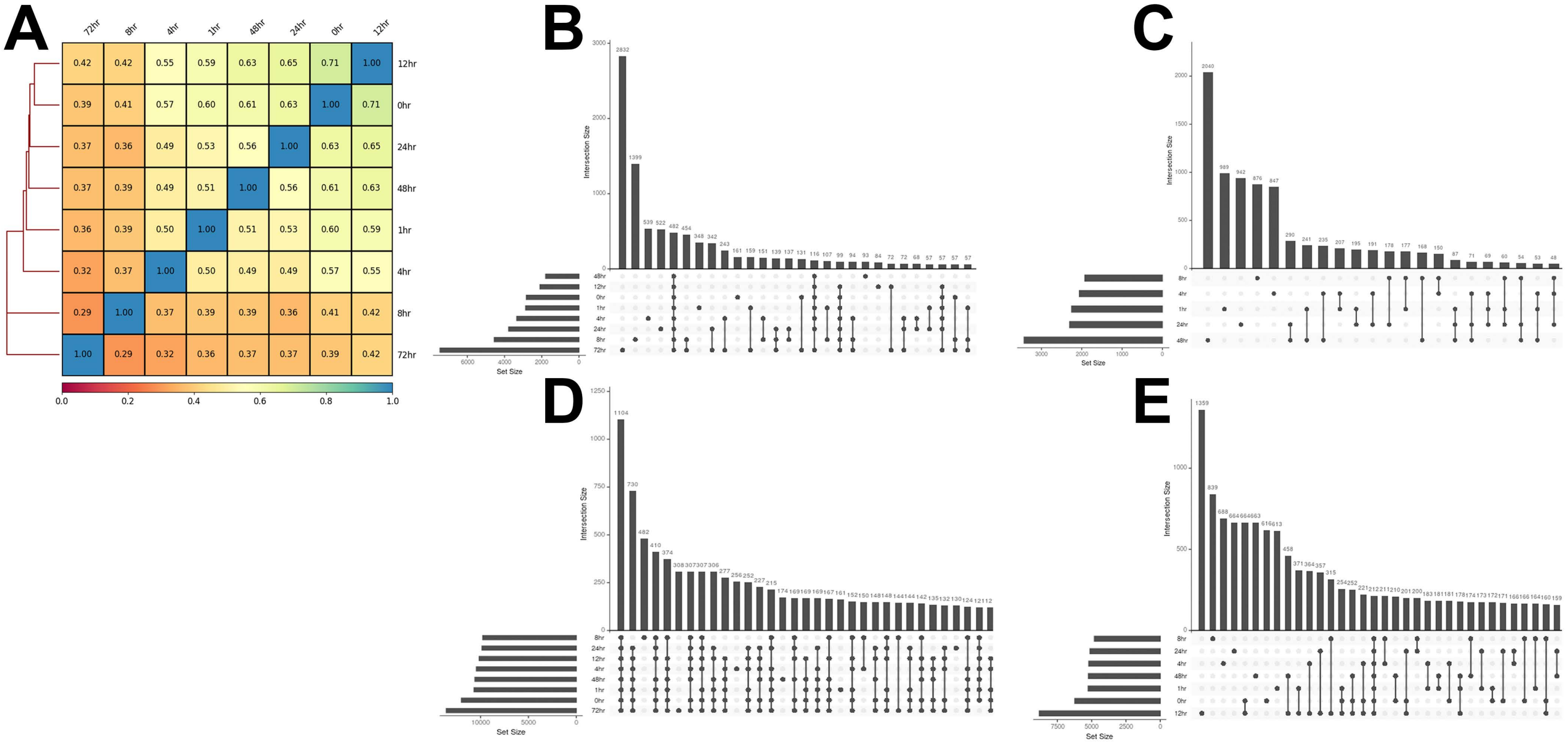

### Supp. Fig. 6. Nucleosome sensitivity is highly variable during the monocyte to macrophage transition

A. Pearson correlation of nucleosome sensitivity in 10bp bins over 2kb promoter regions for all pol II genes. Coefficient values are shown within each square of the heatmap. B. Upset plot showing the number of common and unique genes in the diffusely sensitive cluster (cluster1) for all time points. C. Upset plot showing the number of  shared and unique genes for the strongly positioned sensitive cluster (cluster 2 in 1hr, 4hr, 8hr, 24hr, and 48hr samples). D. Upset plot showing the number of common and unique genes in the strongly positioned resistant clusters (clusters 2-7 for 0hr, clusters 4-7 for 1hr-48hr, and clusters 2-7 for 72hr). E. Upset plot showing the number of shared and unique genes in the mixed clusters (cluster 2 for 1hr, cluster 3 for 1-8hr, clusters 2-3 for 12hr, and cluster 3 for 24-48hr).

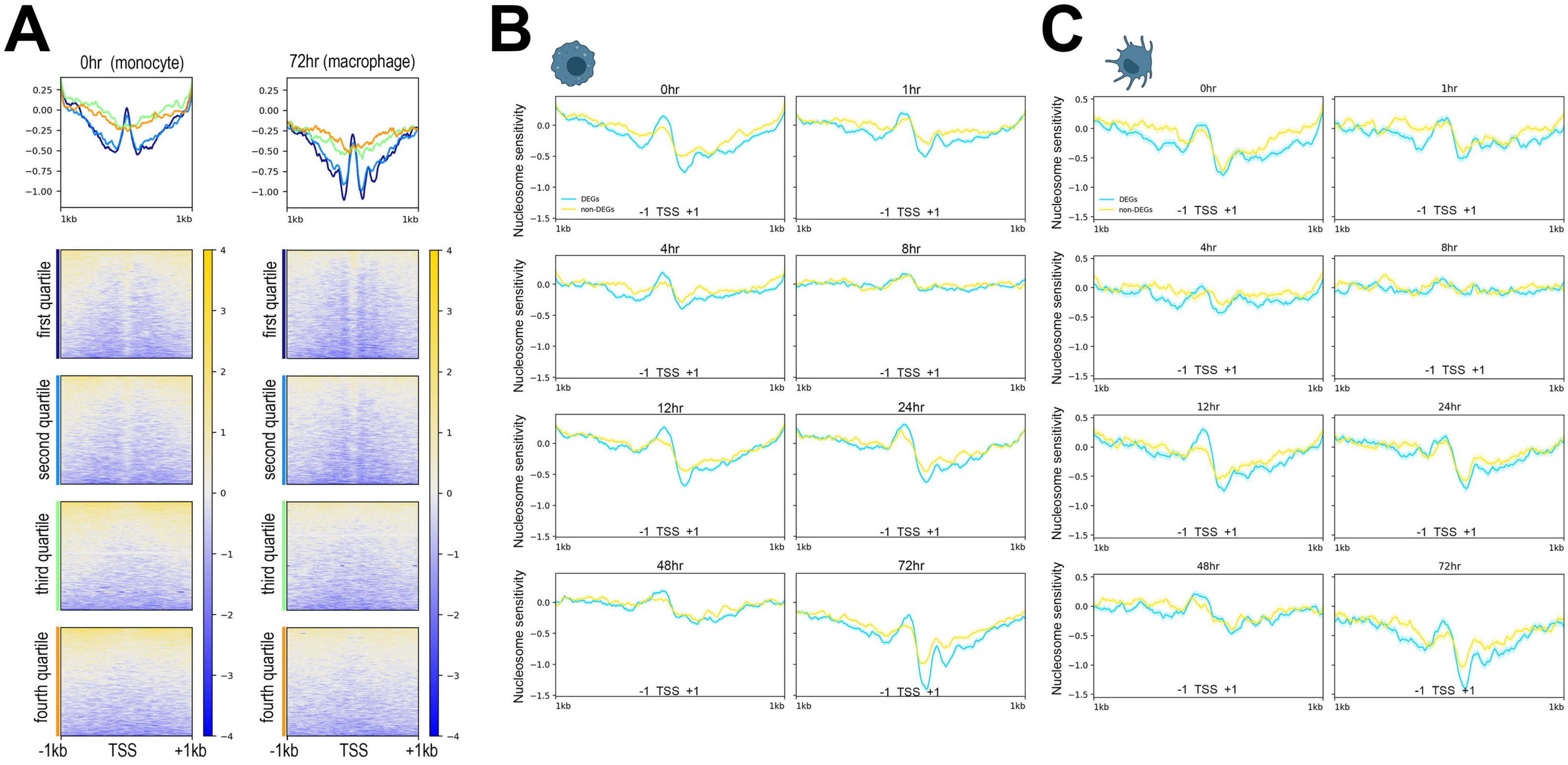

A
0hr (monocyte)
72hr (macrophage)
first quartile
second quartile
third quartile
fourth quartile
-1kb
TSS
+1kb
B
C
Nucleosome sensitivity
DEGs
non-DEGs
-1 TSS +1
0hr
1hr
4hr
8hr
12hr
24hr
48hr
72hr

### Supp. Fig. 7. PMA-responsive and matched promoters experience a contraction in both sensitivity and resistance during PMA-induced monocyte differentiation

A. Heatmaps show sensitivity signal over the TSS of pol II genes separated into quartiles based on gene expression in monocytes (0hr sample) and macrophages (72hr sample). B. Average THP-1 nucleosome sensitivity plotted over PMA-responsive DEGs and a matched set of non-DEG promoters at all time points following PMA treatment. 95% confidence intervals are shown in shaded regions behind average lines. C. Average THP-1 nucleosome sensitivity plotted over monocyte-derived dendritic cells DEGs and a matched set of non-DEG promoters during the monocyte to macrophage transition. 95% confidence intervals are shown in shaded regions behind the average lines

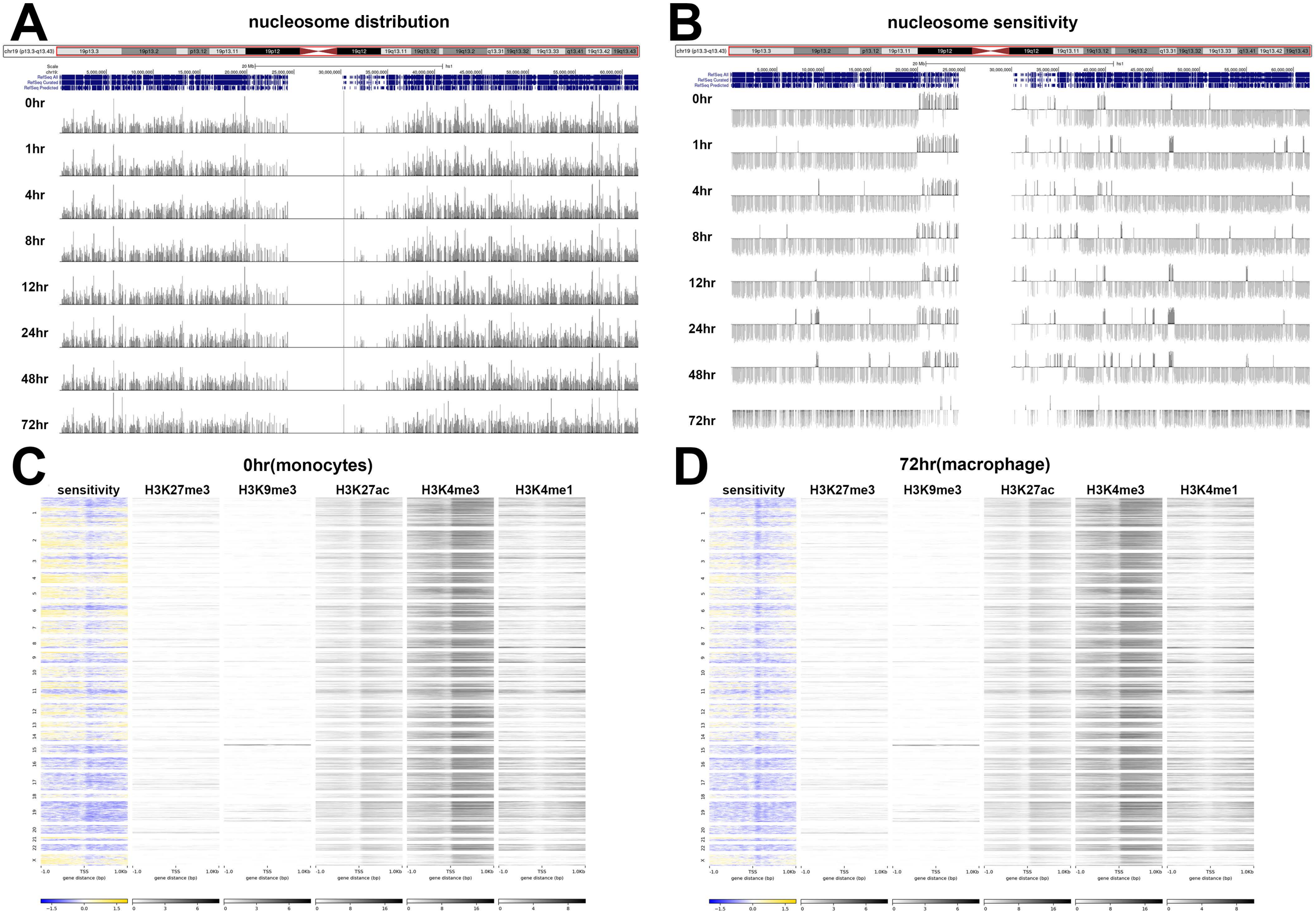
A
nucleosome distribution
0hr
1hr
4hr
8hr
12hr
24hr
48hr
72hr
B
nucleosome sensitivity
0hr
1hr
4hr
8hr
12hr
24hr
48hr
72hr
C
0hr(monocytes)
sensitivity
H3K27me3
H3K9me3
H3K27ac
H3K4me3
H3K4me1
D
72hr(macrophage)
sensitivity
H3K27me3
H3K9me3
H3K27ac
H3K4me3
H3K4me1

### Supp. Fig. 8. Domain-Like Properties of Nucleosome Sensitivity During the Monocyte to Macrophage Transition

A. Genome browser view of nucleosome distribution over chromosome 19 at all time points. B. A genome browser view of sensitivity over chromosome 19 over all time points. Positive values (dark grey) represent sensitive regions and negative values (light grey) represent resistant regions. C. Heatmaps showing nucleosome sensitivity, H3K27me3, H3K9me3, H3K27ac, H3K4me3, and H3K4me1 over promoters in chromosomal order in untreated monocytes. Gene order is identical for all maps. C. Heatmaps showing nucleosome sensitivity, H3K27me3, H3K9me3, H3K27ac, H3K4me3, and H3K4me1 over promoters in chromosomal order in differentiated macrophages. Gene order is identical for all maps.
